# MC1R residue 223 is a convergent target for coat-colour lightening in domestic and wild bovids

**DOI:** 10.64898/2026.10.30.762879

**Authors:** Vitalii Nedosekov, Andrey A. Yurchenko, Jingyi Gao, Aigerim K. Khamzina, Jung-Woo Choi, Nikolay Yudin, Juha Kantanen, Denis M. Larkin

## Abstract

Whether similar phenotypes evolve through shared substitutions or alternative molecular routes is a central question in evolutionary genomics. Grey-coated taurine cattle provide a powerful system for investigating this question using geographically distinct Podolian and Alpine lineages. We combined whole-genome resequencing data, complementary selection scans (hapFLK, DCMS and FST) and contrasts between grey-coated and non-grey-coated breeds to investigate the molecular evolution of grey coat colour. Selection analyses identified candidate regions associated with Podolian/steppe divergence and a prominent pigmentation-related signal on BTA18 contaning *FANCA*, *MC1R*, and *SPIRE2*. Most grey breeds shared a BTA18 haplotype containing MC1R p.Arg223Trp and additional candidate variants. Tyrolean Grey cattle lacked this predominant haplotype but carried the coding duplication p.Gly220_Arg223dup, together with differentiation near the *ASIP* regulatory region. Thus, different candidate coding changes affect the same MC1R residue in different cattle lineages. Haplotype comparisons supported independent Podolian and Alpine origins for p.Arg223Trp and p.Gly220_Arg223dup insertion, respectively. Ancient-genome screening yielded no validated evidence of either variant except p.Arg223Trp in medieval cattle. Comparisons across 1,108 mammalian taxa identified independent recurrence of p.Arg223Trp in the white-coated mountain goat, *Oreamnos americanus*, where the derived allele was fixed in surveyed individuals (JTT-based neutral probability = 0.0043). Our findings support a model in which grey coat colour in cattle evolved through at least two partially distinct molecular routes within interacting pigmentation pathways. More broadly, they identify MC1R residue 223 as a candidate recurrent target of coat-colour lightening and illustrate how domesticated and wild lineages can repeatedly target constrained functional positions during phenotypic evolution.

## Introduction

To what extent adaptive phenotypes could evolve convergently at the nucleotide level in wild animals and domestic breeds remains an underexplored question in evolutionary genomics and domestication research (Cieslak et al. 2011; Stern 2013; Storz 2016). Domestic cattle are a suitable model to address this because of the large number of breeds, their wide geographic distribution from the deserts of Africa to the tundra of Siberia, and remarkable adaptations to diverse environments which together have resulted in substantial genomic diversity since domestication about 10,000 years ago (Decker et al. 2014; Buggiotti et al. 2021; Kim et al. 2023).

Podolian cattle, likely originating from Podolia (modern western Ukraine and northeastern Moldova), represent a group of taurine cattle found in Eastern and Southern Europe and the Mediterranean Basin (Senczuk et al. 2021), making them an attractive model to study adaptive variation. One of the understudied Podolian cattle breeds is the indigenous Ukrainian Grey cattle (UGC), which has historically been shaped both by selection for draught work, meat and milk production, and by local adaptation to the variable steppe environments of Ukraine (Rieznykova 2021). Previous population genetic studies have shown that Podolian cattle, including the UGC, occupy an intermediate position between European taurine and steppe/Turano-Mongolian cattle groups, consistent with a complex demographic history affected by human migrations across Eurasia (Yurchenko et al. 2017).

A distinctive feature of the UGC and many other Podolian cattle is their grey coat colour. Within Podolian cattle, this phenotype varies in shade and intensity and may show age-and sex-related differences (Senczuk et al. 2020; Davidescu et al. 2023). A similar coat colour can be found in other taurine breeds, including the Alpine Tyrolean Grey and the Brown Swiss cattle (Berryere et al. 2003; Senczuk et al. 2020). Studies suggest that coat colour is not only a breed-identifying aesthetic trait but may also be an adaptive phenotype influencing thermal balance, solar radiation absorption, and behavioural responses to environmental stress (Finch et al. 1984; Da Silva et al. 2003; Eizirik and Trindade 2021).

Mammalian coat colour is generally determined by variation in melanocyte development, melanogenesis, pigment-type switching, and pigment distribution within the hair shaft (Reissmann and Ludwig 2013; Baxter et al. 2019; Eizirik and Trindade 2021). In mammals, the *MC1R* and *ASIP* genes play central roles in regulating the balance between eumelanin and pheomelanin production (Klungland et al. 1995; Barsh et al. 2000; Reissmann and Ludwig 2013; Wakamatsu and Ito 2023). Activation of *MC1R* promotes eumelanin synthesis, whereas *ASIP* antagonises *MC1R* signalling and shifts pigmentation towards pheomelanin production (Barsh et al. 2000; Brancalion et al. 2022). Consequently, variants affecting *MC1R*, *ASIP* or interacting pigmentation pathways may produce substantial coat-colour variation. Of particular relevance is the third intracellular loop (IL3) of MC1R, which connects TM5 and TM6 and contains residues critical for G-protein coupling and receptor activation (Wolf Horrell et al. 2016; Ma et al. 2021).

Despite increasing interest in pigmentation genetics, the molecular basis of grey coat colour in taurine cattle remains understudied, particularly at the protein and nucleotide levels. Genome-wide analyses of grey cattle have identified candidate regions on several chromosomes, including BTA (Bos taurus chromosome) 2, BTA4, BTA14, and BTA26, and pointed to genes involved in melanocyte biology, melanogenesis, and pigment transfer (Senczuk et al. 2020). Further work in indicine grey cattle highlighted the complexity of the genetic architecture of this trait (Senczuk et al. 2022). More recently, an analysis of Podolica Italiana grey-coated cattle identified signatures of selection on BTA7 and BTA18, including candidate genes such as *SPATA33*, *SPATA2L*, *FANCA*, and *ZNF276* (Bruno et al. 2024). However, it remains unclear whether different taurine grey-coated breeds share a common molecular basis for coat colour or have evolved similar coat colour phenotypes through different variants.

Coat-colour genes have repeatedly been identified as targets of convergent colour evolution across vertebrates (Mundy 2009), with well-documented examples including independent *MC1R* missense mutations underlying pale coat colour in Gulf Coast beach mice (Hoekstra et al. 2006) and the white “spirit bear” morph of the black bear (Ritland et al. 2001). This demonstrates that the same gene, and sometimes the same functional class of substitutions, can be mutated independently to produce similar colour phenotypes in distantly related species. A question related to this study is whether the mutations controlling grey coat colour in cattle are unique to cattle, are shared with related species expressing a similar coat phenotype or reflect wider nucleotide/amino acid convergence within mammals.

Indeed, studies of another adaptive phenotype, cold adaptation in cattle, show that environmental pressures can result in the same mutations in domestic breeds and wild animals. In Yakut cattle, the world’s northernmost cattle breed, a breed-specific missense variant in the highly conserved *NRAP* gene involved in muscle and heart function was identified as a likely example of convergent evolution associated with adaptation to extreme cold. This variant was absent from other cattle breeds and ancient *Bos* samples, but the same amino acid substitution was observed in multiple cold-adapted mammals (Buggiotti et al. 2021). The *NRAP* example demonstrates that a domestic cattle breed exposed to strong environmental pressures may evolve a molecular solution that parallels one found in distantly related mammals adapted to similar environments. This raises a broader question relevant to taurine grey-coated cattle: whether a coat colour phenotype may similarly evolve through convergent solutions favoured by artificial selection in cattle and natural selection in wild animals.

In this study, we used the UGC together with other Podolian and non-Podolian grey cattle breeds to investigate signatures of selection, identify mutations related to the grey coat colour phenotype, and trace them in >1,000 mammalian taxa to search for convergence events.

## Results

### Placing the UGC cattle within the genetically distinct Podolian cattle group

F*_ST_* analysis between the UGC and 36 taurine breeds based on the SNP genotyping dataset (Yurchenko et al. 2017) revealed a wide range of genetic differentiation, from 0.076 (Kalmyk) to 0.23 (N’Dama) (Supplementary Table S1). Based on these F*_ST_* results, we selected a panel of 21 breeds, of which some were genetically close to, and some were genetically distant from the UGC (Supplementary Table S1).

ADMIXTURE analysis of 22 breeds placed the UGC within the Podolian/steppe cattle group and separated them from Northern European and most other taurine breeds (Figure 1). At K = 6, Podolian breeds formed a distinct cluster, with the shared component substantially present in Kalmyk and Mongolian cattle. This pattern persisted at higher K values (Figure 1). Consistent with ADMIXTURE results, principal component analysis (PCA) placed the UGC close to Kalmyk, Chianina, Mongolian, Podolian, and Romagnola cattle (Supplementary Figure S1). Taken together, these results confirm a distinct genetic component within Podolian cattle and their relationship with other steppe cattle, including Kalmyk and Mongolian breeds.

**Figure 1:**
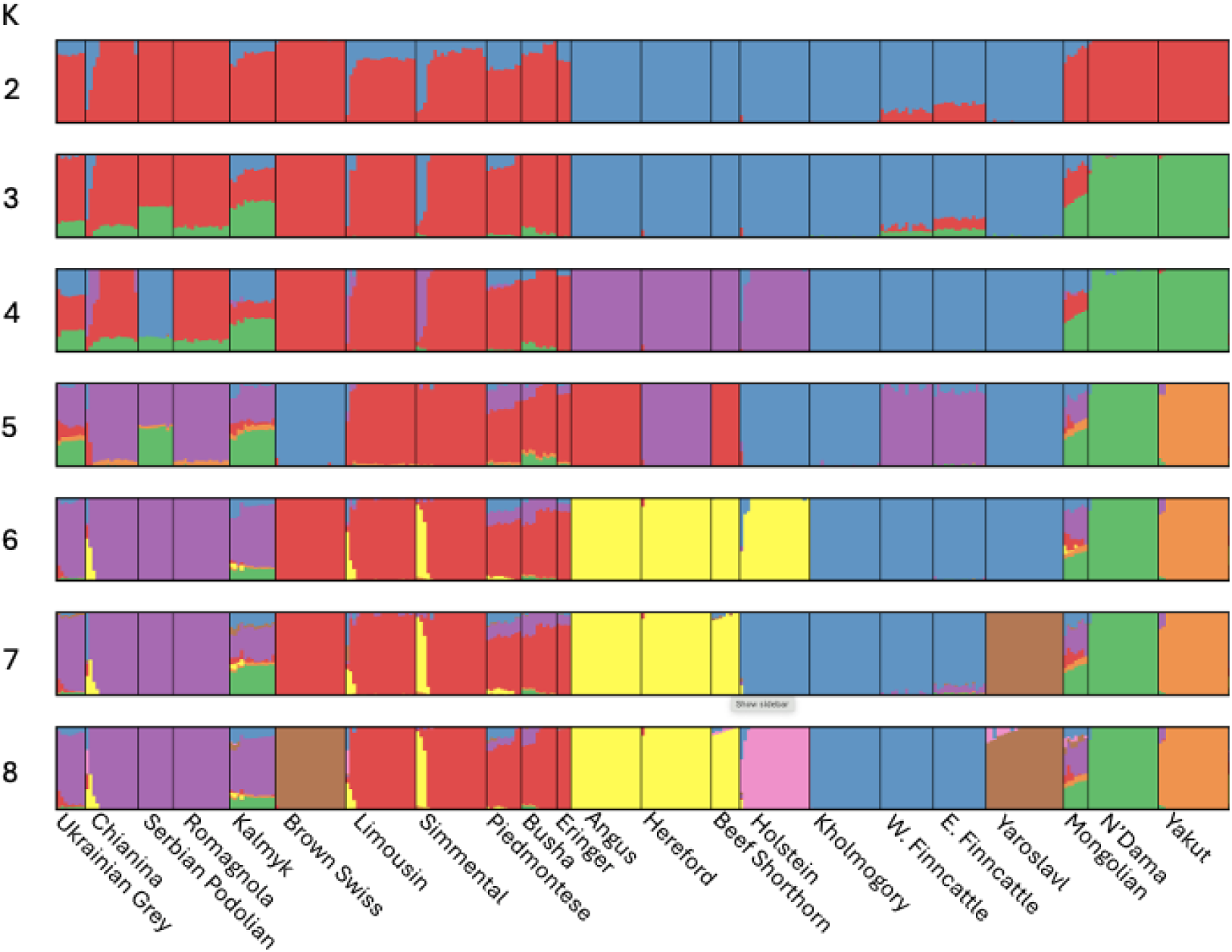
ADMIXTURE analysis of the Ukrainian Grey cattle (UGC) and 21 breeds, both genetically similar to and distant from the UGC, identified through F*_ST_* analysis. K represents the number of assumed populations. W. Finncattle for West Finncattle and E. Finncattle for East Fincattle breeds.

### Scans for signatures of selection

Based on the ADMIXTURE and PCA results we identified two breed groups that we contrasted with the UGC. One group included breeds genetically similar to the UGC: Podolian Chianina and Romagnola, and the steppe Mongolian and Kalmyk cattle (Supplementary Figure S2). The second group included breeds genetically more distant from the UGC: Simmental, Kholmogory, Yaroslavl, Holstein, and Hereford (Supplementary Figure S3). We included the UGC in both groups, with the Yakut cattle as an outgroup where needed. Importantly, the group of closely related breeds included two grey-coated cattle breeds in addition to the UGC (Chianina and Romagnola) whereas the distant breed set contained no grey-coated breeds apart from the UGC.

We then used complementary selection scan methods to identify candidate genomic regions under putative selection in the UGC compared to distant cattle breeds and Podolian/steppe cattle (Supplementary Table S2). In the comparison with genetically distant breeds, hapFLK identified 65 putatively selected regions in the UGC lineage (Supplementary Table S3), while window-based F*_ST_* identified 99 highly differentiated regions in this comparison (F*_ST_* range: 0.53-0.71) (Supplementary Table S4). The genome-wide distributions of hapFLK, window-based F*_ST_*, and point-F*_ST_* are shown in Figure 2.

**Figure 2:**
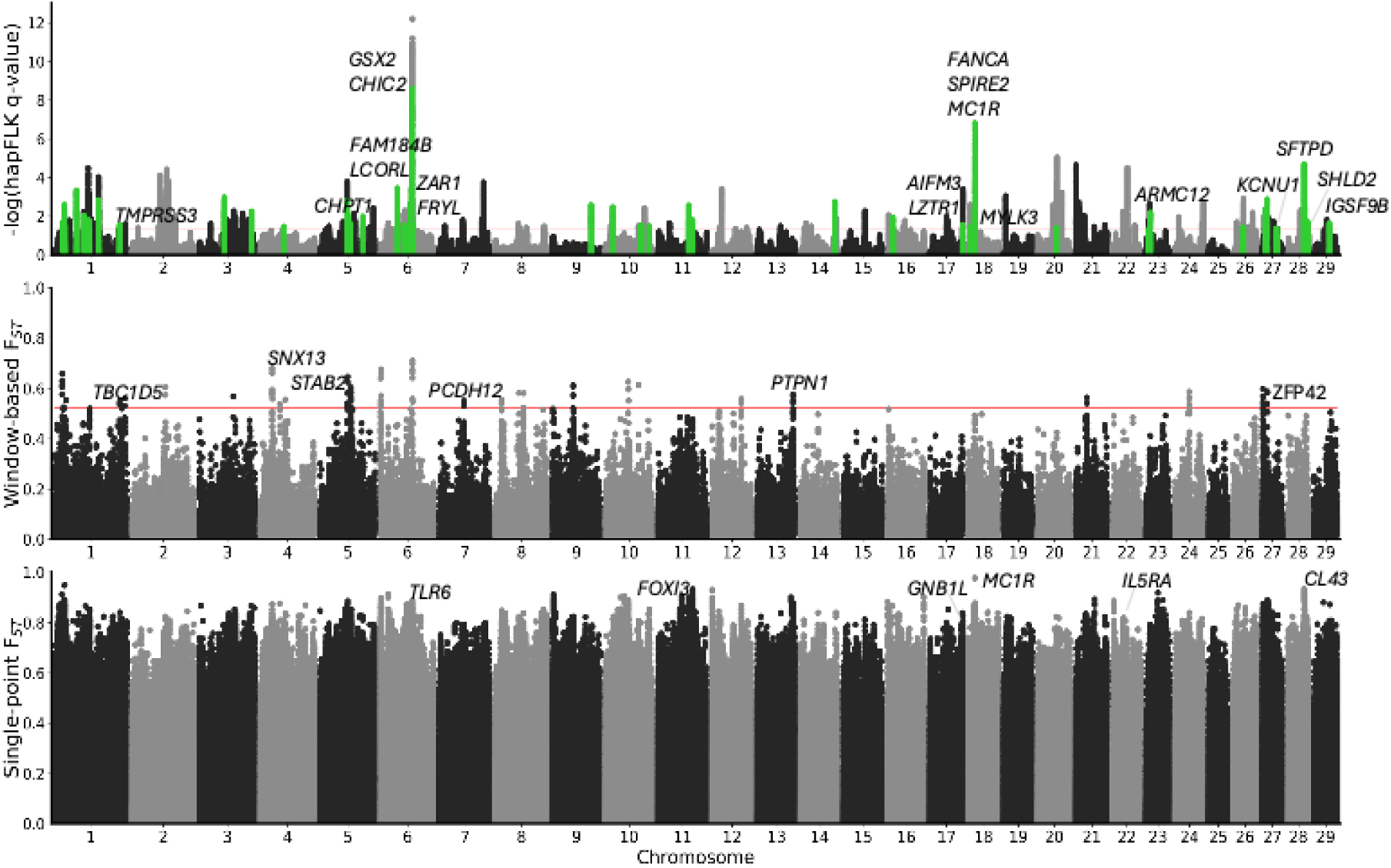
Manhattan plots of hapFLK, window-based F*_ST_*, and point F*_ST_* statistics for the Ukrainian Grey cattle (UGC) samples against a set of genetically distant breeds. The hapFLK plot (green) highlights the selection signatures attributed to the UGC. Red lines denote the q-value = 0.05 threshold for hapFLK and the top 0.1% window value for the window-based F*_ST_* analysis. The top candidate genes are displayed for all three analyses.

The top region in the haplotype analysis contained the genes *CHIC2* and *GSX2*, with *CHIC2* harbouring the top SNP. The second most significant haplotype signal was supported by point-F*_ST_* data and encompassed 23 genes on BTA18, with the strongest signal in *FANCA*. The point F*_ST_* analysis identified a missense mutation in *MC1R*, a coat colour gene neighbouring *FANCA* with F*_ST_* = 0.98, which was the highest signal in the entire genome-wide point F*_ST_* analysis (Figure 2; Supplementary Table S5).

The DCMS analysis of breeds genetically similar to the UGC (Romagnola, Chianina, Mongolian, and Kalmyk) identified 75 regions under putative selection in the UGC, with *SIL1* as the top gene (Supplementary Table S6; Supplementary Figure S4). Similarly, window-based F*_ST_* analysis with these breeds identified *CEBPB* (F*_ST_* = 0.51) as the top gene among the 0.1% of F*_ST_* windows (F*_ST_* range: 0.40-0.51; Supplementary Table S7; Supplementary Figure S4). No coat colour genes were identified in this analysis, including the point-F*_ST_* results, consistent with the absence of substantial differentiation in coat-colour genes in the set containing three grey-coated Podolian cattle breeds (Supplementary Table S8).

### Coat colour genetics

Point-F*_ST_* analysis of the UGC, compared with genetically distant breeds (none of which were grey-coated), identified a missense variant in the key coat colour gene *MC1R* (BTA18:14,706,042) as the top SNP in the entire genome-wide analysis (F*_ST_* = 0.98; Figure 2, Supplementary Table S5). The second most significant selection signal detected in the hapFLK analysis of the UGC, compared with the same breed set, was in the same region, with the top SNP in the gene *FANCA* (BTA18:14,639,215) (Supplementary Table S3).

To test the contribution of the BTA18 region to coat colour in taurine grey breeds, we additionally contrasted grey-coated cattle breeds, as defined by Senczuk and co-workers 2020 (Senczuk et al. 2020) (Chianina, UGC, Tyrolean Grey, Marchigiana, Piedmontese, Romagnola, and Podolian), with non-grey breeds (Angus, Charolais, Limousin, and Holstein) using point-F*_ST_*(Supplementary Table S9). This analysis revealed the strongest F*_ST_* signals on BTA14, followed by signals at several SNPs on BTA18: 14,677,090, 14,609,554, 14,599,929, 14,706,395, 14,706,042, and 14,625,924, which overlapped the haplotype and point-F*_ST_* selection signatures detected in the UGC when contrasted with the distant breed set (Supplementary Table S10). The highest F*_ST_* on BTA14 (0.89) was observed for an SNP in the *PLAG1* gene, whereas the highest F*_ST_* on BTA18 (0.77) was detected for an SNP in the *SPIRE2* gene (Supplementary Table S11).

The frequencies of BTA18 alternate alleles were high in all but one grey-coated breed in the dataset, and in the non-grey-coated breeds, they were zero for most SNPs. The only grey-coated breed in which these alternative alleles were at low frequency or absent was the Alpine Tyrolean Grey. When this breed was excluded from the F*_ST_* comparison between grey- and non-grey-coated breeds, six SNPs on BTA18 showed F*_ST_* values between the grey Podolian and non-grey breed sets ranging from 0.90 to 0.95. The highest F*_ST_* value (0.95) corresponded to an intronic SNP in *SPIRE2*, and two SNPs with F*_ST_* values of 0.94 and 0.93 were in *MC1R*. Two SNPs in the adjacent genes, *FANCA* and *ZNF276*, each had an F*_ST_* values of 0.90. The BTA14 region showed lower F*_ST_* in this comparison than SNPs in *SPIRE2* and *MC1R* (Supplementary Table S11).

Examining allele frequencies in the BTA18 region, we found that one SNP in *SPIRE2*, one in *FANCA,* and two in *MC1R* had an alternative allele frequency of one in the UGC, whereas the other SNPs (in *FANCA* and *ZNF276*) had an alternative allele frequency of 0.56 (Supplementary Table S11). This suggests that SNPs in *SPIRE2*, *FANCA*, or *MC1R* are likely candidate variants underlying the grey coat phenotype. Of these four SNPs, BTA18:14,706,042 in *MC1R* was a missense variant, also identified in the earlier point-F*_ST_* analysis of the UGC against non-grey-coated distant breeds (Supplementary Table S5).

On BTA14, no SNPs had alleles fixed in the grey breed set; however, two SNPs had reference alleles fixed in the non-grey-coated breed set. One of these SNPs was in the 3’ UTR of *LYN*, and the other was intergenic between *LYN* and *RPS20* (Supplementary Table S11). Although the strongest BTA14 signal was associated with *PLAG1*, the presence of grey-coat-associated *PLAG1* alleles in some non-grey-coated animals suggests that this signal may reflect selection on production traits rather than coat colour.

### Independent mutations at MC1R residue 223 as candidates for grey coat colour in cattle

As indicated above, Alpine Tyrolean Grey cattle showed a mutation pattern different from that of the other (Podolian) grey-coated taurine breeds in our comparison. We identified a coding insertion in *MC1R*, starting 17 bp upstream of position 14,706,042, at a frequency of 0.76 in this breed. We scanned animals from the 1000 Bull Genomes Project, Run 9 (Daetwyler et al. 2014; Hayes and Daetwyler 2019), for this insertion and found it in 19 taurine breeds, of which seven carried it at high frequency (alternative allele frequency > 0.6; Supplementary Table S12). Three of these breeds (Tyrolean Grey, Tyrolean Grauvieh, and Maremmana) had grey coat colour, and four (Cabannina, Parthenaise, Tarentaise, and Aubrac) had non-grey coat colour according to the literature (Supplementary Table S12). Of these seven breeds, Maremmana was represented by a single individual, which was homozygous for the duplication, and Cabannina and Parthenaise by two individuals each. Point-F*_ST_* analysis of these two groups of breeds identified the top SNP, BTA13:63,614,182 (FST = 0.90), located ∼20 kb upstream of *ASIP* (Figure 3; Supplementary Table S13).

**Figure 3:**
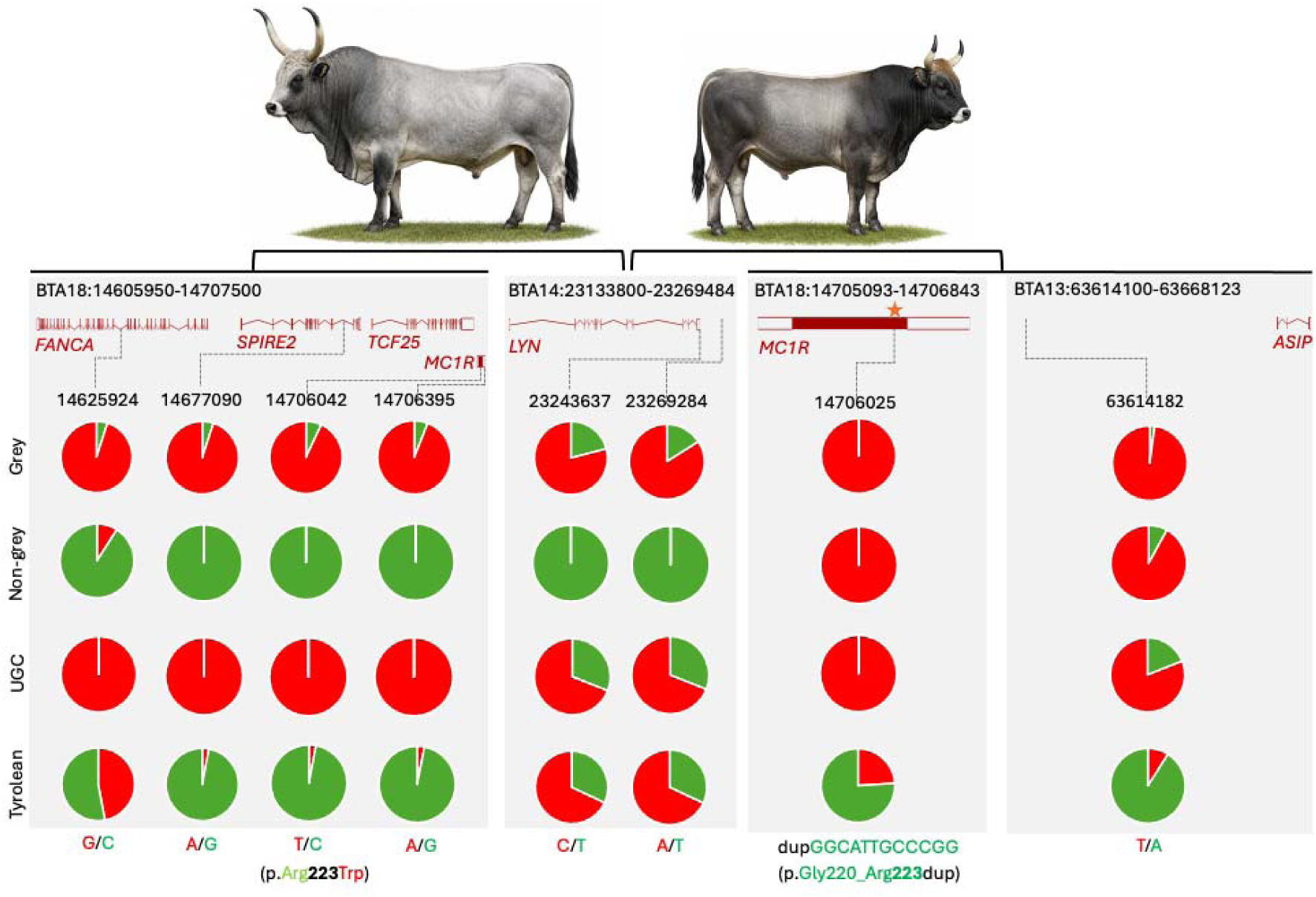
Candidate genomic intervals and allele frequencies for grey coat colour in taurine cattle. The top panel displays the candidate regions and their gene content on BTA13, BTA14, and BTA18. The lower panel shows the positions of the candidate mutations with their allele frequencies listed beneath them. Grey denotes the grey coat-coloured cattle group (all grey-coated breeds except Tyrolean Grey) selected from the 1000 Bull Genomes Project, Run 9. Non-grey-coated breeds are those that do not exhibit grey coat colour. Both sets were selected based on a published classification (Senczuk et al. 2020). UGC, Ukrainian Grey cattle; Tyrolean, Tyrolean Grey cattle. The star marks the location of two genetic variants affecting residue 223 of MC1R.

Interestingly, at the protein level, most Podolian breeds carried an MC1R p.Arg223Trp mutation (BTA18:14,706,042), whereas the Tyrolean cattle carried a p.Gly220_Arg223dup (BTA18:14,706,025). Both mutations are known to affect intracellular loop 3 of MC1R and involve the same residue (position 223) (Kriegesmann et al. 2001; Graphodatskaya et al. 2002; Hauser et al. 2022).

### Distribution of MC1R mutations in the 1000 Bull Genomes Project dataset

We screened 6,191 animals from the 1000 Bull Genomes Project dataset for p.Arg223Trp and p.Gly220_Arg223dup insertion. Both mutations appear likely to be of taurine cattle origin. Among the indicine and hybrid breeds, only one of 16 Beefmaster animals (an indicine-taurine hybrid) carried the p.Arg223Trp variant. For the p.Gly220_Arg223dup, there were four indicine carrier breeds (Butana 11.1%, Ogaden 11.1%, Sheko 4.2%, and Afar 3.6%) and one carrier hybrid breed (Boran 7.5%). Next, we examined both mutations across the taurine breed set. Overall, 18 breeds (n > 7; >14 chromosomes) had the p.Arg223Trp allele. The allele was fixed or nearly fixed in Podolian cattle: Piedmontese (100%), Serbian Podolian (100%), UGC (100%), Romagnola (98.0%), Marchigiana (83.3%), and Chianina (76.7%). Moderate frequencies were observed in Brown Swiss (49.1%) and Kazakh cattle (38.9%). Sporadic occurrence was recorded, among other breeds, in French Alpine (Tarentaise 12.5%), Balkan (Busa 10.0%), Iberian (Limonero 6.3%), Nordic/Baltic (Modern Danish Red 11.1%, Latvian Brown 5.0%, Finnish Ayrshire 4.2%, and Modern Angler 2.5%), and Asian (Buryat 2.5%) breeds (Figure 4).

**Figure 4:**
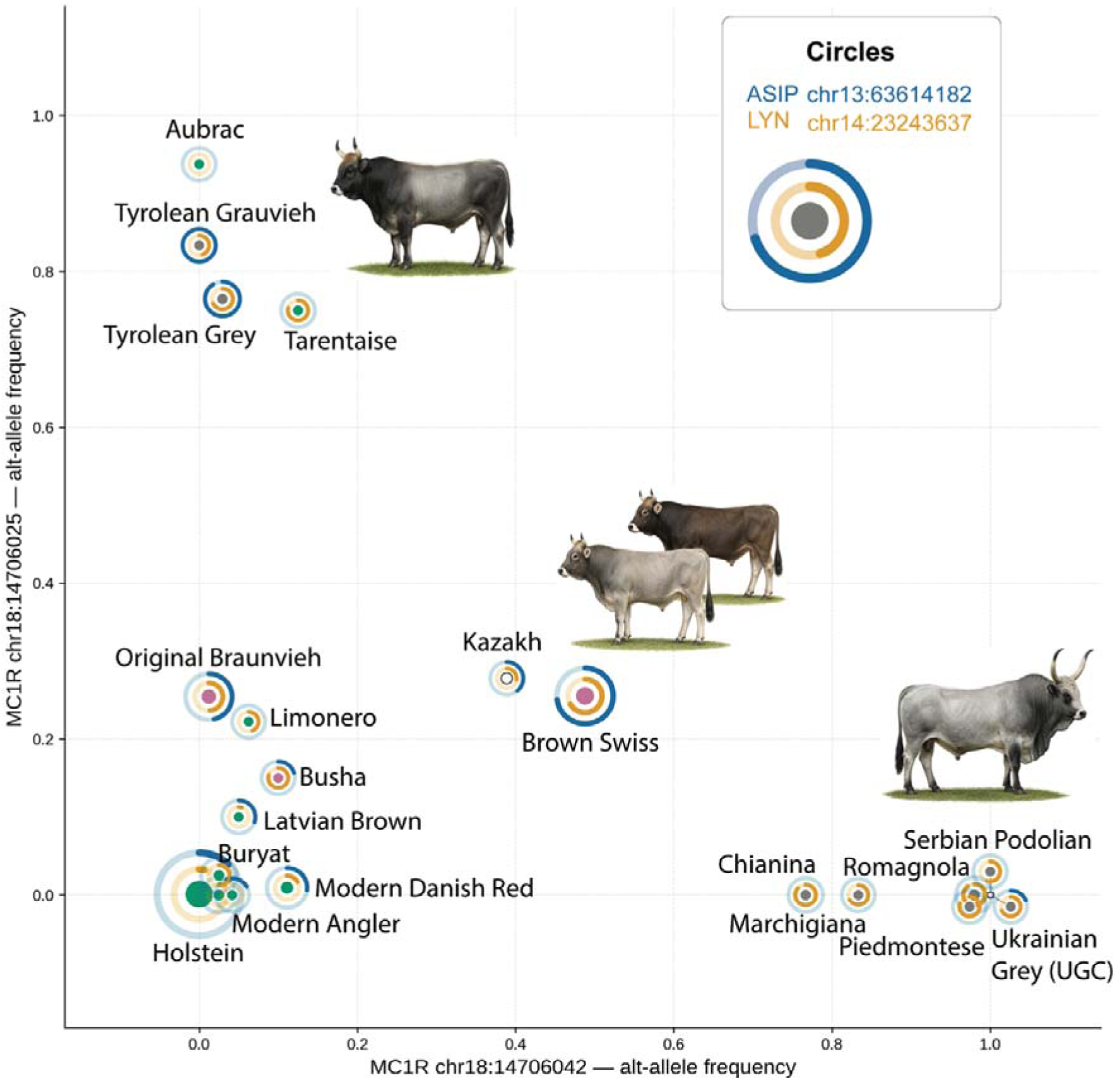
Frequencies of the *MC1R*, *ASIP*, and *LYN* alternative alleles in the 1000 Bull Genomes Project, Run 9, taurine breed set (n > 7 animals per breed; > 14 chromosomes). The X axes show the frequency of the *MC1R* BTA18:14,706,042 missense variant (p.Arg223Trp), while the Y axes show the frequency of the *MC1R* BTA18:14,706,025 coding insertion (p.Gly220_Arg223dup). The outer ring of each circle represents the frequency of the *ASIP* BTA13:63,614,182 variant, and the inner ring shows the frequency of the *LYN* BTA14:23,243,637 variant. The central dot represents breed colour: grey for grey-coated breeds, green for non-grey-coated breeds, white for unknown, and pink for mixed-colour breeds. Dot size represents the number of sequenced animals per breed.

The p.Gly220_Arg223dup was detected across 13 taurine breeds (n > 7). The highest frequencies were observed in Alpine/Swiss and French local breeds: Aubrac (93.8%), Tyrolean Grauvieh (83.3%), Tyrolean Grey (76.5%), Tarentaise (75.0%), Brown Swiss (25.5%), and Original Braunvieh (25.4%). The allele was also present at moderate frequency in Kazakh cattle (27.8%) and Iberian breed Limonero (22.2%), at low frequency, among others, in the Baltic breed Latvian Brown (10.0%), and the Asian breed Buryat (2.5%) (Figure 4). The geographic distribution, centred on the Alpine Arc and extending into French local breeds with additional occurrence in Central Asian and Iberian breeds, differed from the Podolian/Mediterranean distribution of MC1R p.Arg223Trp.

### Origin and dating of the MC1R variants

Pairwise Hamming similarities among the 512 taurine-only subset carrier haplotypes within ±50 kb of the target p.Arg223Trp SNP (3,763 variants) ranged from 95.8% to 100%, with a mean of 99.7%. The haplotype network showed a star-shaped topology with the largest identical group (n = 84; 16.4%), from which 78% of minority haplotypes radiated at distances of 1-3 variants (Supplementary Figure S5). To identify the non-carrier breed most closely related to the ancestral carrier haplotype background, we searched all 9,896 non-carrier taurine haplotypes for the longest stretch identical to the carrier consensus. The longest identical stretch belonged to a Limonero animal, which shared a continuous identical tract of 210.4 kb with the carrier consensus (7,610 variants; 114.9 kb upstream and 95.5 kb downstream of the target SNP), 1.37 times the length of the next-best breed (Brown Swiss, 153.2 kb). When we used only variants distinctive for the carrier haplotype (1,669 informative positions), we detected the longest match (542 variants, 778.5 kb) in Marchigiana, followed by Romagnola (335.4 kb), Limonero (299.1 kb), and Brown Swiss (295.5 kb) (Supplementary Figure S6). Consistent with this, mean pairwise divergence among carrier haplotypes was 1.67-fold higher in Podolian breeds than in Brown Swiss. These results are consistent with a Mediterranean or Podolian cattle origin of the p.Arg223Trp mutation and its introgression into Brown Swiss cattle.

To further test whether the p.Arg223Trp variant entered Alpine Brown Swiss from Podolian cattle, we computed Patterson’s D-statistics (ABBA–BABA; (Patterson et al. 2012) from population allele frequencies, with Original Braunvieh (the ancestral Alpine breed; carrier-allele frequency 1.2%) as P1, Brown Swiss (49.1%) as P2, seven Podolian breeds (Romagnola, Chianina, Serbian Podolian, Piedmontese, Marchigiana, UGC, and Podolica; 92.9%) as P3, and an aurochs individual (homozygous ancestral at the focal site) as the outgroup. The carrier-background *MC1R* haplotype was defined as the variants distinguishing Arg223Trp carrier from non-carrier haplotypes (allele-frequency difference |Δf| ≥ 0.7 between the 510 carrier and 9,898 non-carrier taurine haplotypes; 214 variants spanning 553 kb around the focal site). On this haplotype, D (Original Braunvieh, Brown Swiss; Podolian, aurochs) was strongly positive (D = +0.803, block-jackknife Z = 90.7): Brown Swiss shares substantially more derived alleles with Podolian cattle than Original Braunvieh does, specifically at the *MC1R* carrier haplotype. The excess sharing was localised, across the remainder of the 2 Mb region it disappeared (D = +0.059, Z = 0.59). A control replacing Podolian with Holstein showed no signal (D = −0.032, Z = −0.17), confirming that the localised sharing is specific to Podolian cattle. Results were robust to the carrier-background threshold (|Δf| ≥ 0.5; 463 variants over 1,018 kb: D = +0.732, Holstein D = −0.152, not significant). The direction of gene flow is inferred from the allele-frequency gradient: the mutation is near-fixed in Podolian cattle (92.9%), intermediate in Brown Swiss (49.1%), and essentially absent from Original Braunvieh (1.2%). Together with the shared carrier-haplotype background and the diversity patterns identified above, these results indicate that the mutation was introgressed into Brown Swiss from Podolian cattle.

Pairwise Hamming similarities among the 341 carrier haplotypes within ±50 kb of the p.Gly220_Arg223dup site ranged from 92.7% to 100%, with a mean of 99.7%. The haplotype network showed a star-shaped topology, with the largest identical group (n = 71; 20.8%) from which all minority haplotypes radiated at a median distance of 3 variants. All carrier haplotypes fell within a single cluster (Supplementary Figure S7). To identify the non-carrier breed most closely related to the ancestral carrier haplotype background, we searched all 10,067 non-carrier taurine haplotypes for the longest stretch identical to the carrier consensus. The longest identical tract belonged to a Tarentaise animal (6,602 variants; 183.3 kb), followed by Original Braunvieh (119.6 kb) and Brown Swiss (116.9 kb).

Alpine breeds dominated the upper tail of the tract-length distribution (the 99th percentile was 58.9 kb in Alpine breeds versus 8.2 kb in non-Alpine taurine breeds; the Podolian maximum was 8.2 kb). These results are consistent with an Alpine origin of the p.Gly220_Arg223dup insertion on a Braunvieh-family haplotype (ancestral to Brown Swiss) background, in contrast to the Podolian origin of p.Arg223Trp.

### Distribution of the LYN and ASIP variants

The *LYN* alternative allele (BTA14:23,243,637) was detected across 193 breeds in our dataset. The highest frequencies were observed in Podolian breeds (Romagnola 90.0%, Chianina 86.7%, Piedmontese 80.0%, UGC 68.8%), Alpine/Swiss breeds (Tarentaise 79.2%, Tyrolean Grey 67.6%, Brown Swiss 66.9%), and Simmental-type breeds (Simmental 80.0%, Fleckvieh 77.5%). Notably, the *LYN* alternative allele was also present at high frequency in breeds not associated with grey coat colour (Montbeliarde 61.3%, Jersey 61.1%, Guernsey 45.0%, and Gelbvieh 42.2%) and is prevalent in pure indicine breeds (63.3%). Importantly, an aurochs sample present in the 1000 Bull Genomes Project dataset was heterozygous at the *LYN* site; together with its high frequency across indicine breeds, this suggests that the polymorphism predates domestication and, most parsimoniously, the taurine–indicine split.

Similarly, the *ASIP* alternative allele (BTA13:63,614,182) was detected in 98 breeds, almost exclusively taurine, with an overall allele frequency of 16.5% among taurine cattle; it was nearly absent from indicine cattle (0.17%; two heterozygous carriers among 606 animals) and rare in taurine × indicine crosses (1.6%), suggesting that this is a taurine-specific SNP. The allele was fixed in Tyrolean Grauvieh (100%) and Grey Cattle (100%), and near-fixed in Tyrolean Grey (91.2%). High frequencies were also observed in Brown Swiss (73.0%), Braunvieh (50.0%), and Original Braunvieh (46.4%). Notably, the allele reached a frequency of 80.5% in Jersey cattle, a breed not associated with grey coat colour.

### Screening ancient samples for the MC1R p.Arg223Trp and p.Gly220_Arg223dup variants

We collected 76 aurochs and 120 ancient cattle samples from the Sequence Read Archive (SRA; Supplementarily Table S14). As a prescreen, we queried these samples using tBLASTn (Altschul et al. 1990) with relaxed settings and identified 17 aurochs and 11 cattle samples potentially carrying at least one read consistent with tryptophan at MC1R position 223. We then mapped the reads from all 28 candidate samples to the *MC1R* nucleotide sequence, explicitly guarding against post-mortem C→T deamination, which exactly mimics the C→T substitution underlying the p.Arg223Trp change. Four aurochs candidates had pre-domestication ages (∼46k, ∼41.5k, ∼13k, and ∼10.7k BP). At the nucleotide level, every aurochs candidate with adequate on-target depth, including the two oldest pre-domestication specimens (∼46k and ∼41.5k BP), carried the ancestral C (Arg), and the only T (Trp) observations were isolated reads on a single strand in a post-domestication aurochs (∼9.0k BP) and cattle (∼3.8k BP), a signature of deamination rather than a true heterozygote genotype. The two younger pre-domestication candidates (∼13.0k and ∼10.7k BP) had no recoverable reads at MC1R 223 position and remain unresolved. The single nucleotide-confirmed R/W genotype in the entire dataset was found in the Kaz1 individual (Medieval Caucasus), in which the derived T was supported by multiple interior (non-terminal) reads from independent molecules. We therefore found no nucleotide-validated evidence for the p.Arg223Trp allele in any pre-domestication aurochs (Supplementary Table S14).

Using the same set of ancient samples and tBLASTn conditions, we prescreened for the p.Gly220_Arg223dup peptide and identified 17 aurochs and 16 ancient cattle samples containing at least one read consistent with the presence of the p.Gly220_Arg223dup in MC1R (Supplementary Table S14). We then mapped the reads from all 33 candidate samples to the wild-type and duplication MC1R nucleotide sequences, requiring each supporting molecule to span the duplication junction with at least 5 bp on each side. No ancient samples were positive for the duplication with ≥ 2 independent molecules. One post-domestication cattle sample (∼2.8k BP) had a single read supporting the insertion. We therefore found no nucleotide-validated evidence for the p.Gly220_Arg223dup in any ancient sample, pre- or post-domestication. The summary of breed distribution and origins of the candidate grey coat colour variants is presented in Table 1.

**Table 1.** Origin and dating of *MC1R*, *LYN*, and *ASIP* candidate grey coat colour variants.

| Variant | Modern distribution | Origin | Ancient DNA evidence | Dating |
|---|---|---|---|---|
| MC1R p.Arg223Trp (BTA18:14,706,042) | 4.9% taurine-wide; near-fixed in Podolian breeds (92.9%), intermediate in Brown Swiss (49.1%). | Podolian (Mediterranean); introgressed into Brown Swiss | No validated pre-domestication carrier: all aurochs with genotype calls are ancestral; the sole confirmed carrier is medieval cattle | Post-domestication inferred from modern distribution and ancient sample analysis |
| MC1R p.Gly220_Arg223dup (BTA18:14,706,025) | 3.3% taurine-wide; concentrated in Alpine breeds | Alpine origin | No ancient carrier, pre- or post-domestication | Post-domestication, inferred from modern distribution |
| LYN alternative allele (BTA14:23,243,637) | 193 breeds; 63.3% in indicine cattle | Widespread ancestral polymorphism (no single breed-group origin) | An aurochs from 1000 Bull Genomes Project dataset is heterozygous | Pre-domestication, predates the taurine-indicine split |
| ASIP alternative allele (BTA13:63,614,182) | 98 breeds, almost exclusively taurine | Taurine-specific SNP | No validated ancient carrier (detection constrained by the variant's location within a BovB element) | Pre-/post-domestication dating unresolved |

### Convergent evolution of MC1R residue 223 beyond cattle

To determine whether the same *MC1R* p.Arg223Trp missense variant has appeared in other mammals, we screened the MC1R position 223 in the genomes of 1,116 mammalian taxa from 27 mammalian orders, confirming the MC1R identity in 1,108/1,116 taxa representing 1,034 species and 53 additional subspecies and 21 hybrid or genus-level accessions from 26 orders (133 families) using multi-species protein alignments, tBLASTn against SRA read sets and unannotated WGS assemblies, and BLASTp against the NCBI non-redundant database (Supplementary Table S15; Figure 5). Of these, 874 (78.9%) contained arginine and 228 (20.6%) contained glutamine. Six species (0.54%) contained other amino acids, including one species (0.09%) that contained tryptophan. Within *Cetartiodactyla*, arginine was prevalent in 160/172 taxa, 11/172 taxa contained glutamine (4/7 *Camelidae*, 2/3 *Giraffidae*, 4/81 *Bovidae*, 1/1 *Neobalaenidae*) and one species, *Oreamnos americanus* (Rocky Mountain goat), carried the p.Arg223Trp change (Supplementary Table S15).

**Figure 5:**
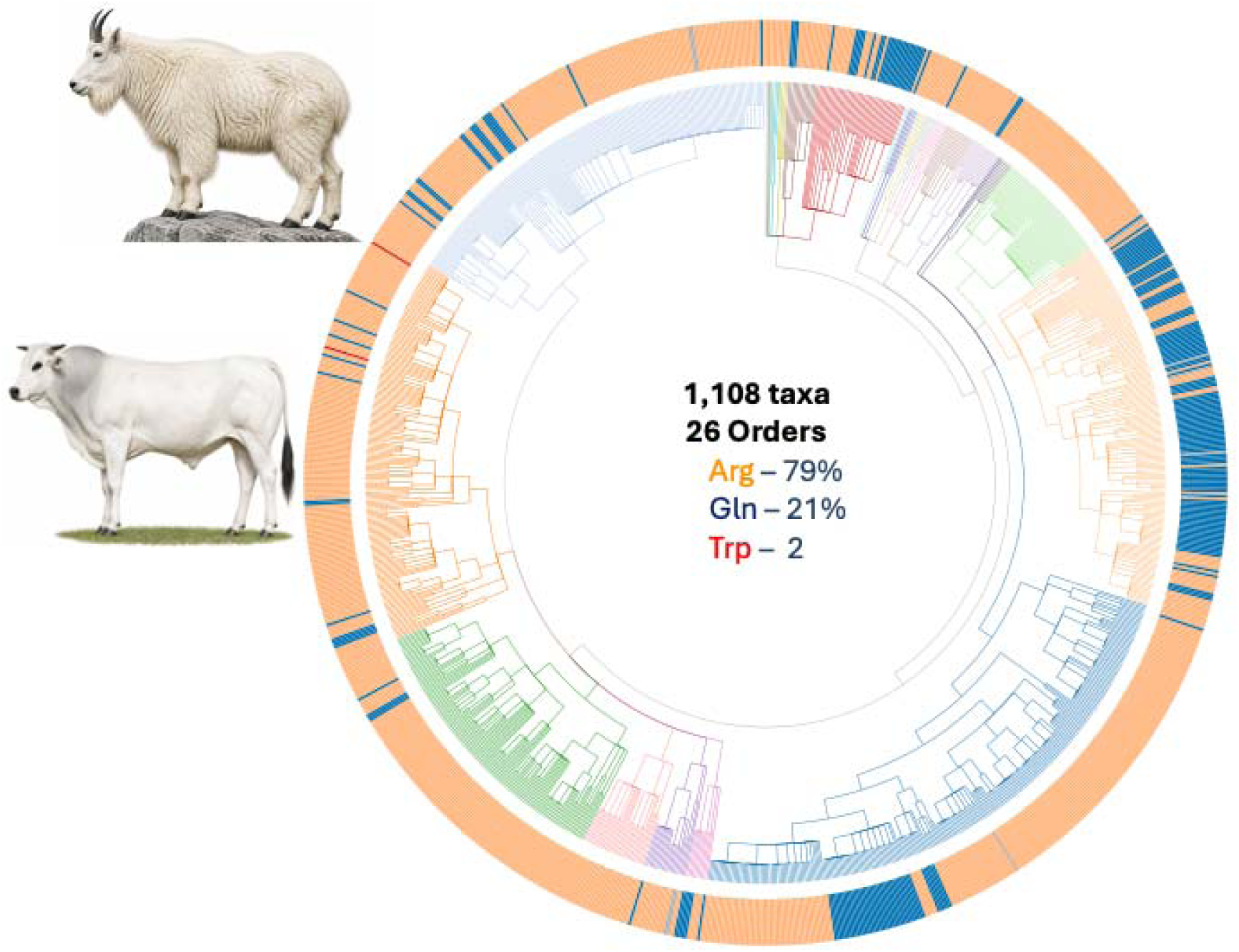
Phylogenetic tree of 1,108 taxa from 26 orders showing the amino acid content at MC1R residue 223. Arginine (Arg) predominates, while glutamine (Gln) is the minor state. Two bovid lineages show tryptophan (Trp): Podolian cattle (represented by the Chianina breed image) and the Rocky Mountain goat. Branch colours indicate different orders.

Strikingly, the coat colour of the Rocky Mountain goat is white, consistent with the *MC1R* p.Arg223Trp change contributing to a pale or reduced-eumelanin coat phenotype in mammals beyond cattle. We examined all 34 *O. americanus* samples available from the NCBI SRA by tBLASTn of a 21-amino acid probe centred at position 223. We confirmed that tryptophan was fixed in all *O. americanus* samples across all sampled geographic locations. No amino acids other than arginine, glutamine, and tryptophan were detected in *Cetartiodactyla* at position 223.

Across mammals, 14 of 26 orders had 100% arginine at position 223; only three orders had glutamine predominating (*Rodentia*, *Diprotodontia*, and *Pholidota*), consistent with independent arginine-to-glutamine substitutions in the therian arginine background. The tryptophan substitutions in *O. americanus* and in Podolian cattle therefore represent two independent derived changes within Bovidae, one at the species level and one at the population (breed) level. *O. americanus* (*Caprinae*) and *Bos taurus* (*Bovinae*) belong to different subfamilies of Bovidae, which is the least inclusive taxonomic family containing both lineages. We therefore restricted the neutral-null test to *Bovidae* (n = 81 taxa in our alignment) on the grounds that a valid same-substitution convergence test requires that both lineages share a similar underlying receptor, ligand, and downstream pigmentation architecture on which the substitution acts. Outside *Bovidae*, cis-regulatory divergence and independent evolution of paralogous pigmentation genes make the phenotypic prediction unreliable even for an identical amino-acid substitution.

We modelled the neutral probability of an arginine (R) to tryptophan (W) substitution using the JTT amino-acid exchangeability matrix (Jones et al. 1992). The JTT exchangeability for R→W is S(R,W) = 126; weighting by the tryptophan background frequency (π_W = 0.0143) and normalising over all arginine substitution rates gives f_JTT(R→W) = 0.0174, meaning that under a neutral JTT null, ∼1.7% of arginine substitutions are expected to be to tryptophan. Within Bovidae, we identified five independent, phylogenetically collapsed species level R→non-R substitution events (four R→Q: *Capra sibirica*, *Ammotragus lervia*, *Madoqua kirkii*, *Neotragus moschatus*; and one R→W: *O. americanus*), with the Podolian cattle R→W representing a sixth, independent occurrence at the breed level. The probability that ≥2 of these six independent events would be R→W by chance is P = 0.0043 (binomial; 0.0051 under a Poisson approximation, λ = 6 × 0.0174 = 0.104). The result is robust to taxonomic scope: expanding the test to the full Cetartiodactyla order (seven independent R→Q events across Bovidae, Giraffidae, Camelidae, and Neobalaenidae, plus two R→W; n = 9 events) yields P = 0.0101 (binomial; 0.0111 Poisson). We note that the Podolian cattle event is a breed-level polymorphism rather than a fixed species-level substitution; restricting the test to species-level events only (*O. americanus* alone, one W among five trials) is not significant (P = 0.084), so the convergence signal resides in the recurrence of R→W across the two levels rather than in either event alone.

## Discussion

Understanding whether convergent phenotypic changes arise through shared or alternative molecular routes remains a central question in evolutionary biology (Stern and Orgogozo 2009; Losos 2011; Stern 2013). Domesticated species provide a particularly informative system for addressing this question, as their phenotypes reflect the combined effects of natural environmental pressures, demographic history, and artificial selection imposed by humans (Andersson and Georges 2004; Andersson 2012; Larson and Burger 2013). In this study, we used grey cattle and wild mammals as complementary models to investigate repeated molecular routes to coat colour change in domestic and wild animals, focusing on the melanocortin-1 receptor (*MC1R*).

Consistent with previous studies (Sermyagin et al. 2018; Senczuk et al. 2021; Solodneva et al. 2023), our population stratification analyses placed Ukrainian Grey cattle (UGC) samples alongside other Podolian cattle. We confirmed a close relationship among Podolian cattle, Nordic cattle, and additional steppe breeds (represented by Kalmyk and Mongolian cattle), supporting a complex demographic history shaped by migration, admixture, and structured breeding across Eurasia (Li et al. 2007; Yurchenko et al. 2017; Davidescu et al. 2023).

Our selection scans, which compared UGC genomes with those of genetically similar Podolian and other steppe breeds, as well as with more genetically distant non-grey-coated breeds, identified genomic intervals and genes under selection in Podolian/steppe cattle (including the UGC) that could contribute to their phenotypes. The comparison with genetically distant breeds highlighted genes potentially associated with the dual-purpose production type of steppe cattle, parasite resistance, climatic adaptations, and coat colour (Rieznykova 2021; Senczuk et al. 2021).

The central evolutionary result of this study concerns grey coat colour. Previous genome-wide studies of grey-coated cattle identified multiple candidate regions involved in pigmentation, melanocyte biology, and pigment transfer (Senczuk et al. 2020; Senczuk et al. 2022; Bruno et al. 2024). Our results extend this work by suggesting that grey coat colour in taurine cattle is unlikely to be explained by a single genetic mechanism. Instead, the data support at least two partly distinct molecular routes involving coding and non-coding variation within interacting pigmentation pathways.

The first route observed in Podolian cattle is associated with the *FANCA–MC1R–SPIRE2* locus on BTA18, where several alternative alleles were fixed or nearly fixed in grey coated breeds and absent or rare in non-grey coated breeds. Within this interval, the *MC1R* missense variant leading to an arginine-to-tryptophan substitution at residue 223 (p.Arg223Trp) demonstrated the strongest F*_ST_* differentiation between the UGC and genetically distant non-grey-coated breeds genome-wide. A linked variant in the 3′ UTR of *MC1R* may also affect expression, although this remains to be tested functionally. The neighbouring *FANCA* and *SPIRE2* signals may indicate additional pigmentation-related or linked effects. *FANCA* has been associated with coat-colour variation in cattle and functions in cellular homeostasis pathways(D’Andrea 2010; He et al. 2022; Kunene et al. 2022), while *SPIRE2* encodes an actin nucleator implicated in organelle dynamics and melanosome transport (Dietrich et al. 2013; Alzahofi et al. 2020).

The second route was observed in Tyrolean Grey and related Alpine breeds. In these breeds, the BTA18 haplotype observed in Podolian cattle was absent or was present at low frequency explainable by later introgression. Instead, Tyrolean Grey carried a coding insertion in *MC1R*, p.Gly220_Arg223dup, previously described as Ed2 (Hauser et al. 2022). Because this insertion was also found in non-grey-coated breeds, it is unlikely to be sufficient on its own to produce grey coat colour. However, when grey-coated and non-grey-coated breeds carrying this insertion at high frequency were contrasted, the strongest differentiation occurred upstream of *ASIP*, a key antagonist of *MC1R* signalling. This region overlaps an extended *ASIP* regulatory interval containing non-coding exons and promoter elements, where regulatory variation has previously been shown to affect coat colour in cattle (Trigo et al. 2021). These results are consistent with a model in which grey coat colour in Tyrolean Grey and other Alpine breeds may arise through a combination of *MC1R* coding duplication and regulatory differentiation in or near *ASIP*.

The BTA14 *PLAG1–LYN* locus may provide an additional shared component across grey-coated cattle. Although *PLAG1* showed strong differentiation, its established role in growth and production traits makes it difficult to attribute the signal specifically to coat colour (Song et al. 2016; Fink et al. 2017). By contrast, variants in or near *LYN* are biologically plausible pigmentation candidates because LYN is a Src-family tyrosine kinase involved in signalling pathways that can influence melanocyte precursor migration and differentiation, including Wnt/β-catenin, NF-κB, and KIT-related signalling (Senczuk et al. 2022; Weerawarna and Richardson 2023). The similar derived allele frequencies in the *PLAG1–LYN* locus in Podolian and Alpine cattle suggest that this region may contribute to both grey colour routes, potentially explaining why BTA14 was repeatedly highlighted in previous studies even when BTA18 signals were not consistently detected, a likely consequence of combining breeds carrying different *MC1R* alleles (Senczuk et al. 2020).

The most striking feature of this two-route model is that candidate MC1R coding changes affect the same amino acid position, residue 223. In the first route, the ancestral arginine is replaced by tryptophan (p.Arg223Trp). In the second route, the same residue is duplicated as part of a short insertion (p.Gly220_Arg223dup). This repeated involvement of the same MC1R position suggests that residue 223 may be involved in coat lightening in taurine cattle and potentially in other species.

Indeed, our results reveal that the identical MC1R p.Arg223Trp substitution arose independently in domestic Podolian cattle and in the Rocky Mountain goat (*Oreamnos americanus*) representing two out of 1,108 mammalian taxa surveyed. In cattle the derived allele segregates within a group of Podolian breeds, reaching fixation or near-fixation in several of them (e.g. UGC, Piedmontese, Serbian Podolian, and Romagnola), while being nearly absent from indicine cattle and from the ancestral Alpine breed Original Braunvieh. In wild *O. americanus* the tryptophan allele is instead fixed across all individuals surveyed. Because cattle and *O. americanus* belong to different subfamilies of Bovidae (Bovinae and Caprinae, respectively) originating from different continents, the two occurrences of this substitution represent genuinely independent origins, one shaped by artificial selection during breed formation and the other by natural selection in a wild population. Against a JTT-based neutral null restricted to Bovidae, the recurrence of this specific change is unlikely by chance (P = 0.0043), supporting genuine molecular convergence rather than a neutral coincidence (Zhang and Kumar 1997).

Both Podolian and *O. americanus* lineages have shifted from the ancestral dark coat toward a pale, non-wild-type phenotype: grey in cattle and white in the Rocky Mountain goat. We therefore interpret residue 223 as a convergent target for coat-colour lightening, while stressing that grey and white may not be the same colours. The functional consequences of ICL3 substitutions in this region are consistent with a role in pigmentation but are not yet resolved for our variants: residue 223 lies adjacent to the K226–R227–Q228–R229 motif shown by alanine-scanning mutagenesis of human MC1R to be essential for G-protein coupling, where substitutions abolish cAMP signalling without affecting ligand binding (Frändberg et al. 1998), and an independent substitution in this same intracellular loop, the goat p.K226E, has been associated with white coat and proposed to be a loss-of-function change (Wu et al. 2006). The only functional characterisation of the cattle p.Arg223Trp allele itself comes from a single in-vitro study in a non-melanocyte cell line (Graphodatskaya et al. 2002), and the *O. americanus* allele has not been tested at all.

This amino acid convergence may carry a wider implication for how new breed phenotypes arise. The p.Arg223Trp allele is not detectable at the nucleotide level in pre-domestication aurochs, with the earliest validated carrier being medieval cattle (Kaz1), consistent with the grey-associated allele being a post-domestication variant rather than pre-existing ancestral standing variation. The p.Gly220_Arg223dup insertion is also likely to be a post-domestication variant. By contrast, the coat-colour-associated *LYN* variant is far older, predating the taurine-indicine split and therefore domestication, representing an ancient standing variation. Breed formation therefore appears to draw on two sources of variation: the recruitment of new, post-domestication variants and the sorting of ancient standing variation and can furnish domestic animals with coat colour features that were not present in their wild ancestors. That artificial selection in cattle and natural selection in a wild bovid have converged on the very same *MC1R* substitution echoes our earlier finding of convergent evolution of cold adaptation between Yakut cattle and wild mammals adapted to cold (Buggiotti et al. 2021). This reinforces the view that human and natural selection can repeatedly arrive at the same molecular solutions providing breeds with features that the ancestral populations did not have but humans desire, e.g. extreme cold adaptation in the Yakut cattle and pale grey colour in the Podolian and Alpine cattle.

In conclusion, our study refines the placement of Ukrainian Grey cattle within the Podolian/steppe taurine lineage and identifies candidate genomic regions associated with adaptive divergence. Most importantly, it suggests that grey coat colour in taurine cattle has evolved through at least two partly distinct molecular routes involving coding and regulatory changes in interacting pigmentation pathways. The repeated involvement of MC1R residue 223 implicates this site as a potential evolutionary hotspot in coat colour lightening in bovids. Together with the *NRAP* example in Yakut cattle, these findings support the broader view that domesticated cattle can evolve adaptive phenotypes similar to those of wild animals through recurrent changes in constrained functional genes, shaped by the combined action of natural and artificial selection.

## Materials and Methods

### Samples and data

Blood samples were collected from twenty UGC individuals at the Altai Experimental Agricultural Farm in Cherga village, Shebalinsky District, Altai Republic. Sampling was performed during routine veterinary examinations by a licensed veterinarian using standard jugular venepuncture into EDTA vacutainer tubes. The procedure was approved by the Ethical Committee of the Institute of Cytology and Genetics, Siberian Branch of the Russian Academy of Sciences (approval 37/28, November 2017). Informed consent was obtained from the farm owner to use residual biological material for DNA extraction and genomic analyses. No experimental interventions beyond standard veterinary practice were performed. Genotyping of these UGC individuals was performed using the Illumina BovineSNP50 DNA Analysis BeadChip. Genotype data were processed in PLINK v2.00 (Purcell et al. 2007; Chang et al. 2015) to assess pairwise relatedness among animals (--genome). Ten unrelated genotyped individuals with pairwise PI_HAT < 0.125 and no common sires or dams, as determined from pedigrees, were selected for whole-genome sequencing on the Illumina HiSeq 4000 platform (Novogene, Cambridge, UK), generating over 50 Gb (∼16x mean coverage) of paired-end reads per sample. Additionally, ten previously sequenced UGC genomes were obtained from the 1000 Bull Genomes Project, Run 9 (Supplementary Table 2).

Sequence data were processed following the 1000 Bull Genomes Project protocol (Daetwyler et al. 2014; Hayes and Daetwyler 2019). Adapter trimming and quality filtering were performed in Trimmomatic v0.38 (Bolger et al. 2014) in paired-end mode using the following settings: LEADING:20, TRAILING:20, SLIDINGWINDOW:3:15, AVGQUAL:20, and MINLEN:35. Clean reads were aligned to the bovine reference genome (ARS-UCD1.2_Btau5.0.1Y) using BWA-MEM v0.7.17 (Li 2013). Duplicate reads were marked with Picard v2.18.2 (https://broadinstitute.github.io/picard/) using OPTICAL_DUPLICATE_PIXEL_DISTANCE=2500 to account for the sequencing platform. Base quality score recalibration (using the ARS1.2PlusY_BQSR_v3 dataset) and variant calling were performed with GATK v3.8-1-0-gf15c1c3ef (McKenna et al. 2010; Depristo et al. 2011; Van der Auwera et al. 2013). Following previously published protocols (e.g. Igoshin et al. 2024), only single-nucleotide polymorphisms (SNPs) were retained for downstream analyses. Variants were subjected to stringent quality control using the GATK VariantFiltration tool with the following hard-filtering thresholds: *QD < 2.0*, *FS > 60.0*, *MQ < 40.0*, *MQRankSum < −12.5*, and *ReadPosRankSum < −8.0*, as recommended by GATK Best Practices (Van der Auwera et al. 2013).

### Dataset creation

To identify breeds that are genetically similar to or distant from the UGC, we first combined the 20 genotyped UGC individuals with taurine breeds previously genotyped (Yurchenko et al. 2017) (sample size > 10) and calculated Hudson’s F*_ST_* in PLINK v2.00 (Supplementary Table S1). The distant-breed contrast was designed to identify broader lineage-specific divergence, whereas the close-breed contrast was intended to filter out the shared Podolian/steppe background and prioritise signals more specific to the UGC.

#### Population analysis dataset

Autosomal SNP data from whole-genome resequencing for 22 selected breeds were extracted from the 1000 Bull Genomes Project, Run 9 VCF files. ADMIXTURE v1.3 software (Alexander et al. 2009a) and principal component analysis (PCA) implemented in the SNPRelate R package (Zheng et al. 2012) were used to identify four breeds genetically closest to the UGC in the 1000 Bull Genomes Project dataset (Romagnola, Chianina, Mongolian, and Kalmyk), and five genetically distant breeds (Hereford, Holstein, Kholmogory, Simmental, and Yaroslavl). Yakut cattle samples were included in the distant-breed analyses as an outgroup where needed. This dataset was used solely to identify breeds genetically closest and distant to the UGC using a high-density SNP set.

#### Signatures of selection datasets

Raw sequencing data for these breeds were downloaded from the NCBI Sequence Read Archive (Supplementary Table 2) and processed using the same mapping and SNP detection pipeline described in the *Samples and data* section. Two variant datasets were created: (a) UGC and the four genetically similar breeds, and (b) UGC and the five genetically distant breeds plus Yakut cattle (as an outgroup). fastSTRUCTURE (Raj et al. 2014) was used to detect and exclude admixed individuals from these sets (Supplementary Figures S2 and S3). The final two datasets comprised of (a) UGC (n=20), Romagnola (n=19), Chianina (n=20), Mongolian (n=20), Kalmyk (n=20), and (b) UGC (n=20), Hereford (n=20), Holstein (n=20), Kholmogory (n=20), Simmental (n=20), Yaroslavl (n=20), and Yakut (n=20) (Supplementary Table 2).

#### Coat colour dataset

To identify candidate mutations associated with grey coat colour, a dataset was extracted from the 1000 Bull Genomes Project, Run 9. It included Chianina (n=15), Ukrainian Grey (n=10), Tyrolean Grey (n=17), Marchigiana (n=9), Piedmontese (n=10), Romagnola (n=25), and Serbian Podolian (n=10) in the grey-coated cattle group, and Angus (n=20), Charolais (n=20), Limousin (n=20), and Holstein (n=20) in the non-grey-coated cattle group. Breed classification followed Senczuk and coworkers (2020). This dataset was used solely to contrast grey-and non-grey-coated breeds.

### Scans for signatures of selection

To detect genomic regions under selection in the UGC population, we applied three complementary methods: hapFLK (Fariello et al. 2013), the de-correlated composite of multiple signals (DCMS) (Ma et al. 2015; Verity et al. 2017), and the fixation index (F*_ST_*)(Weir and Cockerham 1984). The hapFLK analysis was performed on a dataset of breeds genetically distant from UGC, whereas the DCMS approach was applied to a dataset of breeds genetically closest to UGC. Multiallelic variants were decomposed into biallelic sites, and the resulting high-quality SNP datasets were used for the hapFLK and DCMS analyses. The final datasets for the genetically closest and distant breed groups contained 38,405,479 and 31,103,195 SNPs, respectively.

For hapFLK, additional filtering was performed in PLINK v2.00 to exclude rare variants (*--maf 0.1*) and variants with >10% missing genotypes (*--geno 0.1*). No further filtering beyond the initial hard-filtering step was applied to the DCMS dataset. The F*_ST_* analyses were conducted on both the genetically closest and distant breed groups without minor-allele-frequency filtering. All analyses were performed on autosomes only.

#### HapFLK analysis

To investigate breed-specific signatures of selection, we applied hapFLK v1.4 using a workflow adapted from an earlier study (Igoshin et al. 2024) and optimised for the UGC dataset. First, we estimated the number of haplotype clusters by running fastPHASE v1.2 (Scheet and Stephens 2006) on a panel including UGC and genetically distant breeds, yielding *K* = 30. HapFLK v1.4 (Fariello et al. 2013) was then run with *K =* 30 and the --nfit=30 option, and cluster-frequency estimates were written to files using the --kfrq flag.

We summarised statistical significance by fitting a robust linear model (rlm function, MASS R package (Venables and Ripley 2002) and deriving *P*-values from the resulting normal approximation. q-values were estimated with the qvalue R package (Storey and Tibshirani 2003). A genomic region was considered to show evidence of selection if it contained at least one SNP with a q-value < 0.01. Region boundaries were defined by the first flanking markers on either side with q-values > 0.2. For each identified region, we reconstructed phylogenies with *local_reynolds.py* (https://forge-dga.jouy.inra.fr/attachments/3248) and *local_trees.R* (https://forge-dga.jouy.inra.fr/attachments/10561) to determine which populations showed the signal, and we examined local haplotype composition using the hapflk-clusterplot.R script (https://forge-dga.jouy.inra.fr/attachments/2919) to validate these observations.

#### DCMS analysis

As outlined in our previous study (Igoshin et al. 2024), the DCMS approach combines several correlated selection statistics into a single de-correlated composite statistic to increase overall sensitivity (Ma et al. 2015). In this study, the DCMS analysis incorporated the H1, H12 (Garud and Rosenberg 2015), iSAFE (Akbari et al. 2018), XP-nSL (Szpiech et al. 2021) and T (Harris and DeGiorgio 2020) statistics. Prior to these analyses, the VCF datasets were phased using SHAPEIT v4.2 (Delaneau et al. 2019), assuming an effective population size of 1,000. The Markov Chain Monte Carlo parameters were set to “-- mcmc-iterations 10b,1p,1b,1p,1b,1p,1b,1p,10m”. Recombination rates were estimated using a likelihood-based genetic map (Qanbari and Wittenburg 2020).

The H1 and H12 statistics were computed using the script *H12_H2H1.py* (https://github.com/ngarud/SelectionHapStats/blob/master/scripts/H12_H2H1.py) with the following parameters: window size = 200 SNPs, step = 10 SNPs, and maximum number of mismatched alleles = 4 (Garud and Rosenberg 2015). The iSAFE method (Akbari et al. 2018) was run using *isafe.py* v1.0.7 (https://github.com/alek0991/iSAFE/releases/tag/v1.0.7). Calculations were performed in 3 Mb windows with 1 Mb increments, and results from all intervals were merged after excluding the first and last 1 Mb of each window to avoid edge effects. The XP-nSL analysis was conducted with *selscan* v1.3.0 (Szpiech and Hernandez 2014; Szpiech et al. 2021). The T statistic was computed with *lassip* v1.1.0 (DeGiorgio and Szpiech 2022) Haplotype spectra were first generated using the options *--calc-spec --hapstats --lassi* in sliding windows of 200 SNPs with steps of 10 SNPs, followed by the calculation of T statistics for these windows (*--spectra --lassi*). Output statistics from all methods were combined into a single BED file using BEDTools (Quinlan and Hall 2010). Covariance matrices among the five statistics were estimated with the “CovNAMcd” function from the rrcovNA R package (Todorov et al. 2011), with alpha =0.75 (fraction of SNPs sampled) and nsamp = 500,000 (number of random samples). Final DCMS scores for each SNP were calculated with the “DCMS” function in the MINOTAUR R package(Verity et al. 2017). Significance thresholds and boundaries for selected regions were determined as described in the hapFLK section.

#### F_ST_ analysis

Single-nucleotide polymorphisms and genomic regions showing elevated differentiation between the target and reference cattle populations may underlie breed-specific traits. To identify such loci, we calculated F*_ST_* between UGC and pools of genetically closest and genetically distant breeds. VCFtools v0.1.13 (Danecek et al. 2011) was used for the Weir and Cockerham’s F*_ST_* analyses(Weir and Cockerham 1984). Window-based F*_ST_* values were computed using the parameters *--fst-window-size 50000*, *--fst-window-step 25000*, and *--max-missing 0.9*. For point-F*_ST_* estimation, only the window parameters were omitted. Based on the window-based analysis, genomic intervals with F_ST_ values in the top 0.1% of windows (excluding those with F_ST_ < 0.4) were considered outliers, following the criteria of Igoshin et al. 2024.

#### Identification of the candidate mutations for the grey coat phenotype

To identify SNPs with high F*_ST_* between grey-coated and non-grey-coated cattle breeds, we used animal genotypes extracted from the 1000 Bull Genomes Project (see *Dataset creation: Coat colour dataset*). We performed Weir and Cockerham’s point-F*_ST_* analysis between these groups in VCFtools v0.1.13. The allele frequencies of the top F*_ST_* variants were calculated for the grey-coated and non-grey-coated groups.

A manual inspection of the Tyrolean Grey animals’ BAM files (IGV; (Robinson et al. 2011)) revealed a 12-bp insertion at BTA18:14,706,025. All animals in the unfiltered 1000 Bull Genomes Project, Run 9 variant calls (including indels) were screened for this insertion. Breeds with an insertion frequency > 0.6 were classified as grey-coated or non-grey-coated based on breed descriptions (Anon 2016) and publicly available images, and point-F*_ST_* was calculated in VCFtools as described above.

### Origin and dating of the MC1R variants

#### Panel and phasing

Analyses used the BTA18 ±1 Mb region around *MC1R* (92,557 biallelic variants (SNPs and indels) from the 1000 Bull Genomes Project dataset, Run 9, ARS-UCD1.2). Because both focal alleles are largely restricted to cattle, haplotype analyses used a pure-taurine subset (Species annotation "Taurus"; n = 5,204 of 6,191 samples), re-phased *de novo*, not filtered from full-panel phasing, with SHAPEIT5 v1.0.0 (phase_common, -- region 18 --thread 8 --seed 1, cattle linkage map (Qanbari and Wittenburg 2020). p.Arg223Trp carriers were defined by the derived allele at the focal SNP BTA18:14,706,042 (345 carriers; 512 carrier haplotypes).

#### Haplotype similarity and network

Pairwise Hamming distances were computed among carrier haplotypes over the ±50 kb window around each focal variant (3,763 variants for p.Arg223Trp; 3,764 for p.Gly220_Arg223dup; the two focal sites are separated by only 17 bp, so the two windows are offset by 17 bp and differ by a single variant). Carrier haplotypes were defined by the derived allele at each focal site separately (512 and 361 carrier haplotypes, respectively). Unique haplotypes were enumerated over the same window, and the haplotype network was constructed as a minimum spanning tree (SciPy;(Virtanen et al. 2020)) on pairwise Hamming distances between unique haplotypes, visualised in NetworkX (Hagberg et al. 2008) using the unweighted Kamada–Kawai layout. Radiation was quantified as the Hamming distance from each minor unique haplotype (all unique haplotypes except the predominant one) to the predominant haplotype.

#### Ancestral haplotype background

For each focal variant, the carrier consensus haplotype was defined as the majority allele among carrier haplotypes at each variant across the ± 1 Mb region (92,557 biallelic variants). For each non-carrier taurine haplotype the longest contiguous stretch identical to the carrier consensus around the focal site was computed, and the haplotype with the longest stretch was retained for each breed. For the carrier-distinctive analysis, informative positions were defined as variants whose carrier-consensus allele had a frequency < 0.5 among non-carrier taurine haplotypes (1,669 positions for p.Arg223Trp; 1,510 for p.Gly220_Arg223dup), and stretches were extended across informative positions only. Mean pairwise Hamming distances among carrier haplotypes were computed within breed groups over the ±50 kb window. Podolian breeds comprised Romagnola, Chianina, Serbian Podolian, Marchigiana, Piedmontese, UGC, Maremmana, and Podolica.

### Introgression of the p.Arg223Trp into Brown Swiss cattle

#### Panel and populations

Analyses used the full SHAPEIT5-phased panel: 6,191 samples (12,382 haplotypes) genotyped at 92,557 biallelic variants within the *MC1R* locus (BTA18:13,705,164–15,706,830, ARS-UCD1.2), phased with SHAPEIT5 v1.0.0 as described above. Four populations were defined from the panel metadata: P1 = Original Braunvieh (n = 125), P2 = Brown Swiss (n = 294), P3 = the seven Podolian breeds (Romagnola, Chianina, Serbian Podolian, Piedmontese, Marchigiana, Ukrainian Grey, Podolica; n = 78), and as outgroup a single aurochs individual (Bos primigenius; SAMN04028906), which is homozygous ancestral (C/C) at the Arg223Trp focal SNP (18:14,706,042). Holstein (n = 1,148) served as a negative-control P3. All Original Braunvieh, Brown Swiss, Podolian and Holstein samples had Species == "Taurus" in the panel metadata. Derived-allele frequencies at the focal SNP were computed from genotype counts: Original Braunvieh 1.2% (3/250 alleles), Brown Swiss 49.1% (289/588), Podolian 92.9% (145/156).

#### Carrier-background SNP set

Carrier haplotypes were defined as taurine haplotypes carrying the derived allele at the Arg223Trp focal marker within the full-panel SHARPEIT5 phasing (510 carrier vs 9,898 non-carrier taurine haplotypes). For each of 92,557 variants in the approximately 2 Mb region we computed |Δf|, the absolute allele-frequency difference between carrier and non-carrier taurine haplotypes. The carrier-background set was defined as variants with |Δf| ≥ 0.7, yielding 214 variants spanning 553 kb around the focal site (BTA18:14,153,577–14,706,395). Robustness was assessed with a relaxed threshold (|Δf| ≥ 0.5: 463 variants spanning 1,018 kb; BTA18:13,793,498–14,811,507). For each threshold, the "outside" control set comprised all remaining variants in the region.

#### D-statistic computation

Patterson’s D (ABBA–BABA; (Patterson et al. 2012)) was computed from population allele frequencies using the ratio-of-sums estimator: D = Σ□(p_₂_□ − p_₁_ □)(p_₃□_− p_O□) / Σ (p_₁□_+ p_₂□_− 2p_₁□_ p_₂□_)(p_₃□_+ p_O□ − 2p_₃□_p_O□), where p₁□, p_₂□_, p_₃□_ and p_O□ are the derived-allele frequencies in P1, P2, P3 and the outgroup at variant i. Frequencies were calculated by counting ALT alleles, including both alleles of the diploid aurochs individual. We used the ABBA-minus-BABA convention, so positive D indicates excess sharing between Brown Swiss (P2) and Podolian cattle (P3) relative to Original Braunvieh (P1). Standard errors were estimated using an unweighted delete-one-block jackknife over ten consecutive, near-equal-variant-count blocks ordered by genomic position. Blocks were constructed separately for each selected variant set and its complement. We reported Z D/SE D.

### Ancient sample analysis

We collected ancient Bos genomes from the NCBI SRA/ENA (199 samples: 78 aurochs, 121 cattle). After excluding samples flagged as modern, libraries annotated as RNA-seq or amplicon/metagenomic the final screening set comprised 196 ancient samples (76 aurochs, 120 cattle; Supplementary Table S14). Each sample’s sequencing reads were screened by tBLASTn using short MC1R peptide probes corresponding to the *Bos taurus* MC1R protein sequence encoded by NM_174108.2 at four window sizes (9, 13, 21, and 31 aa): probes centred on residue 223 in both ancestral-Arg (Arg223) and derived-Trp (Trp223) versions, and probes centered on the novel copy1|copy2 junction of the 12-bp duplication allele (c.658_669dup; p.Gly220_Arg223dup), with the 223 Arg and Trp probes used as wild-type junction controls. To accommodate short, damaged ancient-DNA reads we applied relaxed hit thresholds (BLAST E-value 1.0; hits reported at ≥ 50% amino-acid identity over ≥ 8 aligned residues), and a read was counted as duplication-supporting only if its alignment spanned the junction with more identical residues than the best achievable wild-type cross-match (7/9, 9/13, 17/21, and 27/31 residues for the four probe lengths). The union of the codon-223 and duplication candidate-follow-up lists comprised 52 samples (26 aurochs and 26 cattle), represented by 863 unique sequencing runs. Both variants were evaluated in every sample using the same validation pipeline. The modern positive control ERR4836512 was analysed separately and excluded from ancient-cohort totals. Reads were obtained as original ENA FASTQs or through validated full-quality SRA extraction using SRA Toolkit v3.4.1 (Leinonen et al. 2011). Sequencing adapters were removed using Cutadapt v5.2(Martin 2011). All retained reads were aligned using BWA aln v0.7.19 with −l 1024 -n 0.03 -o 2, followed by bwa samse (Li and Durbin 2009). Reads were aligned to six *MC1R* capture sequences: two without the duplication, carrying C or T at c.667, and four with the duplication. Captured reads were then aligned separately, using the same BWA parameters, against two complete bovine genomic references: ARS-UCD1.2 (GCF_002263795.1), including all 2,211 contigs, and an otherwise identical reference containing the verified 12-bp *MC1R* duplication. The corresponding wild-type and duplication-bearing coding sequences were 954 and 966 bp long. BAM processing used samtools v1.22.1 (Danecek et al. 2021) and read-level evidence was extracted using pysam v0.23.3 (https://github.com/pysam-developers/pysam). Genomic specificity required a best alignment at *MC1R* with mapping quality ≥20. Reads with an equally good or better non-*MC1R* primary alignment or alternative alignment reported in the BWA XA tag were excluded. We compared each read’s alignment to the genomes with and without the duplication using edit distance, which counts mismatches and inserted or deleted bases. Corresponding MC1R positions were tracked between the references, and observations at c.667 were excluded when their alignment or assignment to a duplicated copy was ambiguous. We also verified that alignment to either genome preserved the original captured-read sequence and complete base-quality scores.

Evidence was collapsed conservatively within each sample before variant assessment. Mates sharing a sequencing-run/spot identifier were combined into a single allele vote, and observations sharing mapped endpoints and orientation were linked across runs, using paired outer endpoints where available. Discordant allele observations within a family were excluded from support for that variant.

For c.667C>T, the diagnostic base was required to have base quality ≥20. After family collapse, an observation was classified as damage-safe when the diagnostic base lay at least five bases inward from each trimmed read end, or, for a terminal observation, when the same allele was represented on both strands. Supported derived-T evidence additionally required at least two damage-safe T-supporting families. Ancestral-C support was recorded separately.

For c.658_669dup, a read supported the duplication only if it matched the duplication reference with fewer edits than the nonduplicated reference and aligned continuously across the new junction, covering at least five bases on each side. All ten junction-flanking bases were required to have base quality ≥20. Nonduplication support was assessed using the same criteria but requiring a better match to the nonduplicated reference. Reads matching both references equally well were not counted as allele-specific support. Supported duplication required at least two qualifying endpoint-defined families; a single duplication-supporting family remained unresolved. The C→T terminal-damage safeguard was not applied to the insertion itself. Evidence for c.667C>T in reads aligned to the duplication-bearing reference was evaluated using the same base-quality, read-position and family-support criteria described above.

#### Scanning mammalian genomes for MC1R protein position 223

We applied a three-level search for MC1R protein position 223 in mammalian genomes available from the NCBI Genome and SRA databases. First, a multi-species protein alignment was performed on server NCBI using COBALT with default parameters (Papadopoulos and Agarwala 2007). A 21-amino-acid motif centred at bovine MC1R protein position 223 was extracted for each species from the alignment. Second, this dataset of motifs was used to perform tBLASTn (Camacho et al. 2009) searches against unannotated genomes on NCBI. For each species, motifs from the same order or phylogenetically closest orders were used. Additional 21-amino-acid motifs were extracted and combined with the first set of motifs. Lastly, this combined set of motifs was used for tBLASTn searches against SRA sequencing runs of additional mammalian species, using motifs from the same or phylogenetically closest orders for each species. The resulting motifs were collected and the final set of motifs was used for blastp against NCBI protein non-redundant (nr) database.

Species with MC1R top hits from BLASTp alignments (E-value < 0.05; ≥ 75% identity) were retained in the final dataset to avoid matches to paralogous melanocortin receptors.

#### Neutral-null convergence test at MC1R position 223

The neutral probability that an arginine substitution produces tryptophan was estimated from the JTT amino-acid exchangeability matrix ((Jones et al. 1992); machine-readable matrix jones.dat from PAML,(Yang 2007)). The conditional probability was computed as f_JTT(R→W) = S(R,W)·π_W / Σ_{j≠R} S(R,j)·π_j, where S(R,W) = 126 is the JTT exchangeability between arginine and tryptophan, π_W = 0.014261 is the JTT background frequency of tryptophan, and the denominator (103.256) sums the rates of all arginine-to-non-arginine substitutions. This gives f_JTT(R→W) = 0.0174; that is, ∼1.7% of neutral arginine substitutions are expected to be to tryptophan.

Independent substitution events at position 223 were enumerated from the species alignment by collapsing taxa that share a derived residue through common ancestry (NCBI taxonomy tree); for example, glutamine in *Lama* and *Vicugna* (*Lamini*) and in the two Giraffa species were each counted as single events. The breed-level p.Arg223Trp polymorphism in Podolian cattle was counted as one additional independent event, because the convergence hypothesis spans species-and population-level substitutions. All non-arginine calls were verified against source records (NCBI protein accessions, or tBLASTn of the cattle MC1R protein against genome assemblies). Under the null, each independent R→non-R event lands on tryptophan with probability p = 0.0174. The probability of observing ≥2 R→W events among n independent events was computed as the binomial tail P(X≥2) = 1 − (1−p)^n − n·p·(1−p)^{n−1} and cross-checked with a Poisson approximation (λ = n·p) in Python (scipy.stats). The test was performed within Bovidae (n = 6 events: four R→Q, two R→W) and, as a sensitivity analysis, across Cetartiodactyla (n = 9 events: seven R→Q, two R→W). The taxonomy tree used for event collapsing is a proxy containing polytomies; the single polytomy affecting event counts (Camelidae) was resolved using the established sister relationship of Lama and Vicugna.

## Data availability

All raw sequencing data generated in this study for Ukrainian Grey cattle animals are available from NCBI SRA under the BioProject accession number PRJNA1347415.

## Acknowledgements

The authors thank the 1000 Bull Genomes Project group for providing the data on world cattle breeds for our analyses.

## Supplementary Material

Supplementary data are available at *Molecular Biology and Evolution* online

## Author contributions

Vitalii Nedosekov: Formal analysis; Investigation; Visualisation; Writing-original draft; Writing-review & editing.

Andrey Yurchenko: Formal analysis; Methodology; Writing-review & editing. Jingyi Gao: Formal Analysis.

Aigerim Khamzina: Formal Analysis.

Visualisation; Writing-original draft; Writing-review & editing.

Jung-Woo Choi: Investigation; Resources; Writing-review & editing.

Nikolay Yudin: Supervision; Writing-review & editing.

Juha Kantanen: Investigation; Resources; Writing-review & editing.

Denis M. Larkin: Conceptualisation; Methodology; Project administration; Supervision; Writing-review & editing.

