## Supplemental figures for "MC1R residue 223 is a convergent target for coat-colour lightening in domestic and wild bovids": Grey_cattle_suppl_figures_final.pptx

### Slide 1
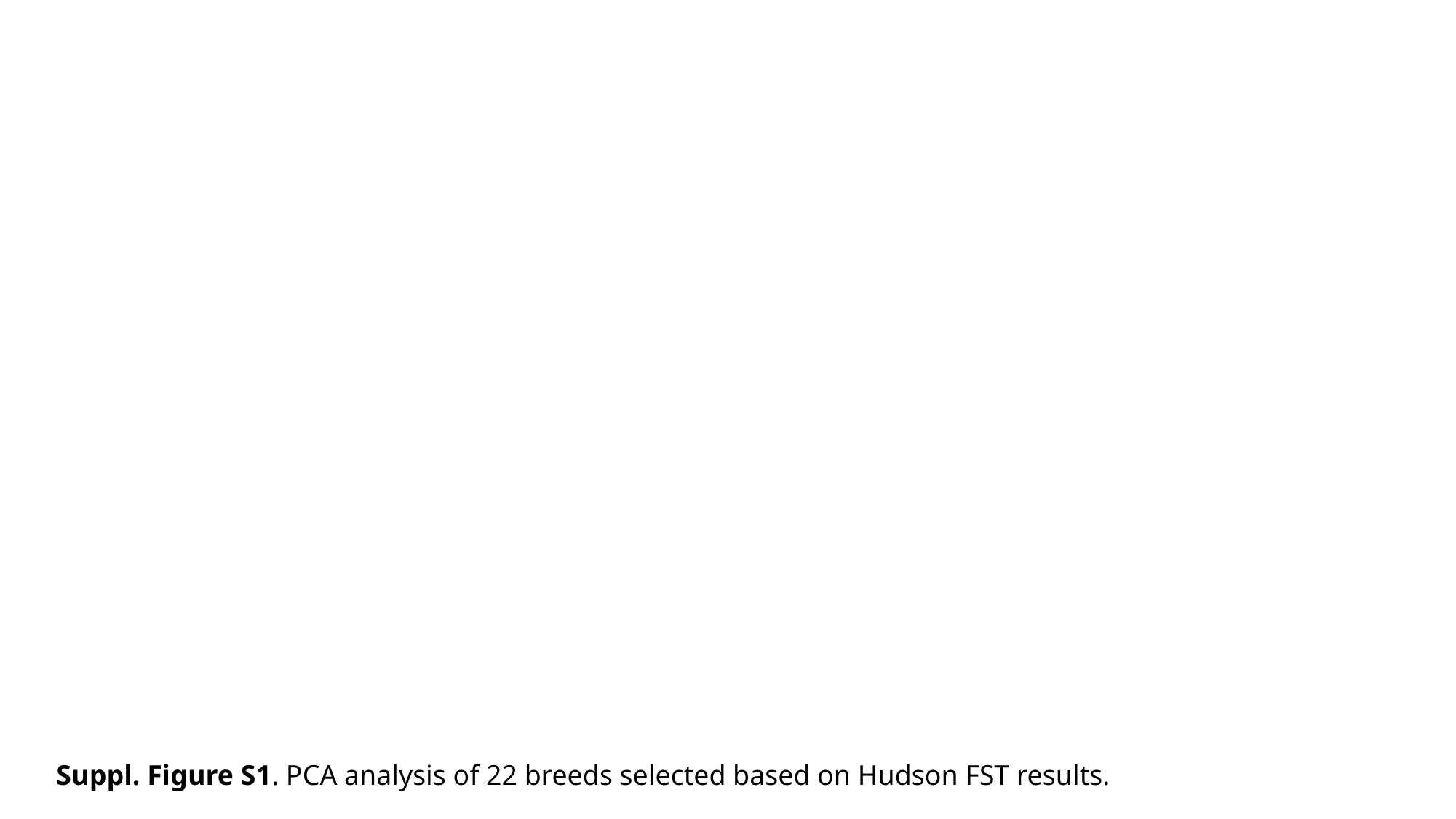

Suppl. Figure S1. PCA analysis of 22 breeds selected based on Hudson FST results.

### Slide 2
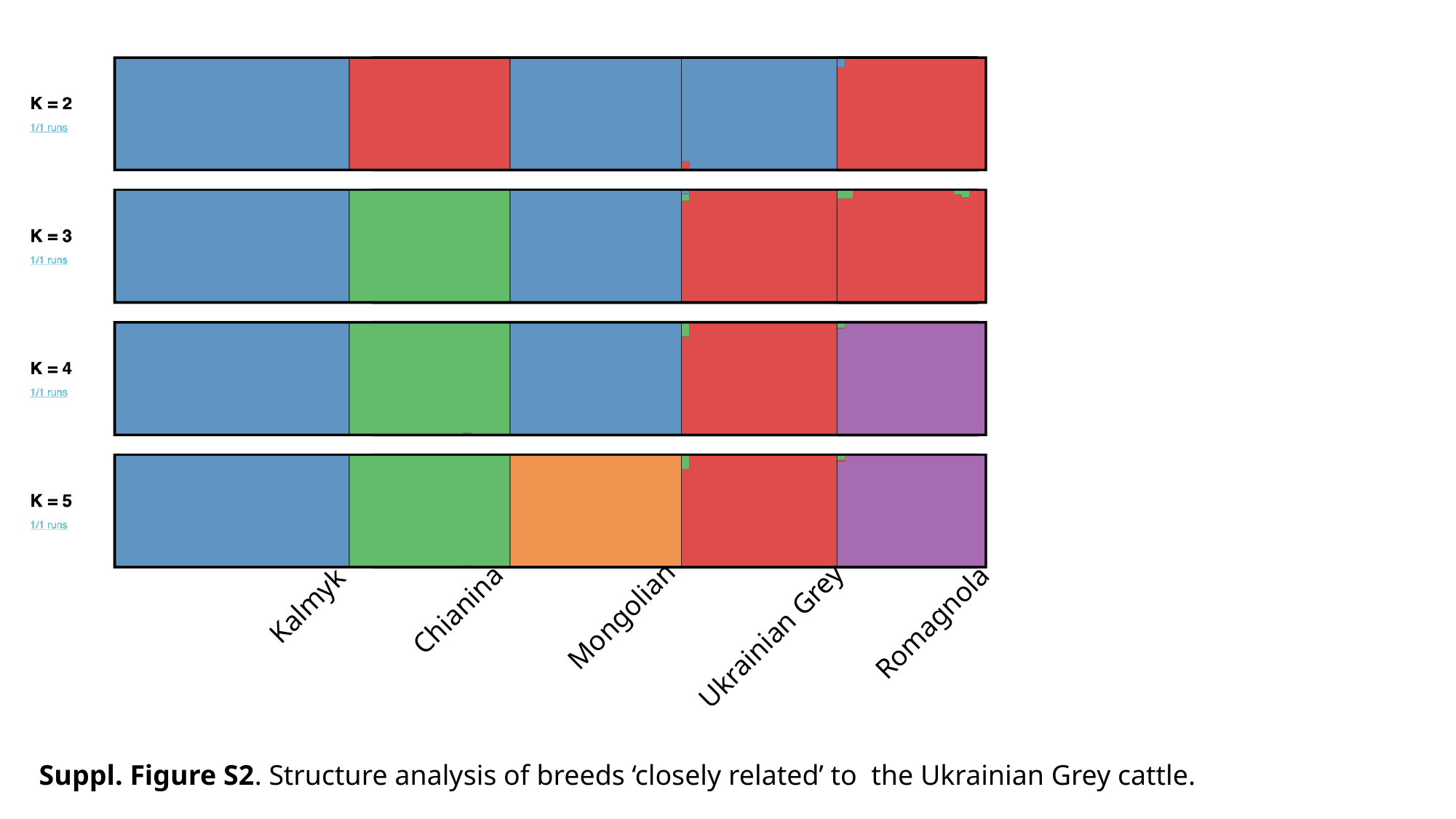

Kalmyk
Chianina
Mongolian
Romagnola
Ukrainian Grey
Suppl. Figure S2. Structure analysis of breeds ‘closely related’ to the Ukrainian Grey cattle.

### Slide 3
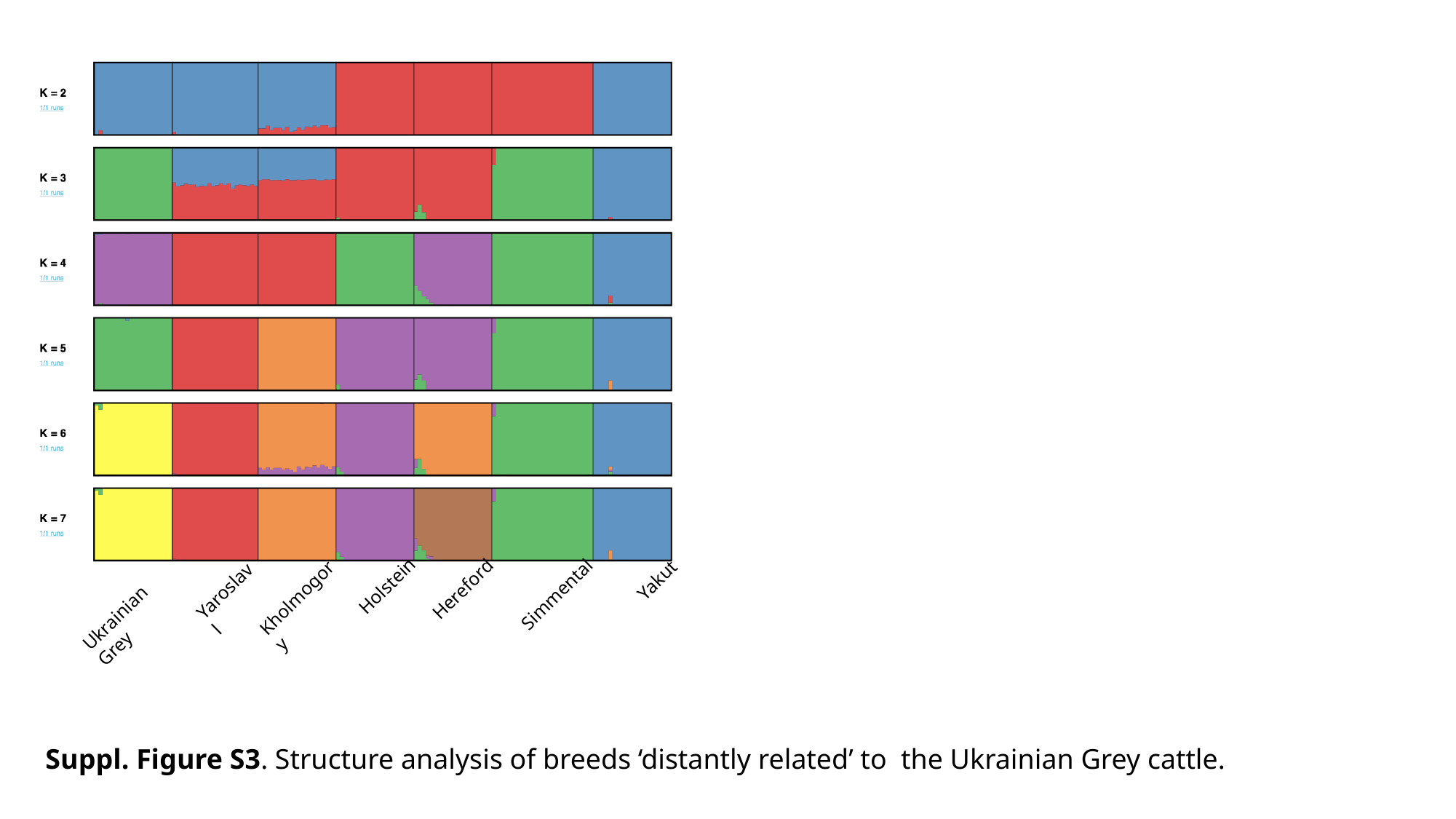

Yakut
Holstein
Hereford
Yaroslavl
Simmental
Kholmogory
Ukrainian Grey
Suppl. Figure S3. Structure analysis of breeds ‘distantly related’ to the Ukrainian Grey cattle.

### Slide 4
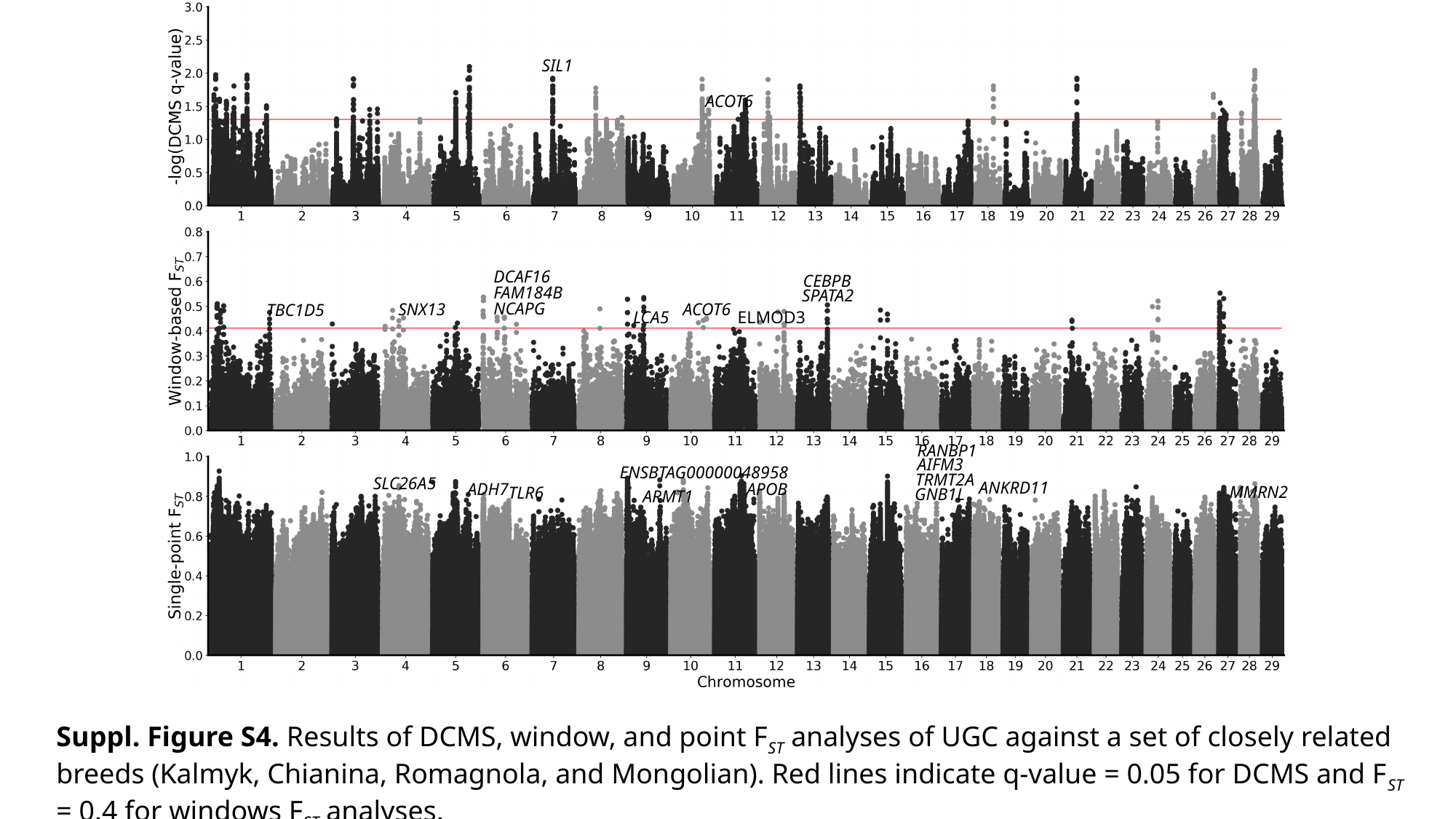

SIL1
ACOT6
DCAF16
CEBPB
FAM184B
SPATA2
NCAPG
SNX13
ACOT6
TBC1D5
LCA5
ELMOD3
RANBP1
AIFM3
ENSBTAG00000048958
TRMT2A
SLC26A5
ANKRD11
ADH7
APOB
MMRN2
TLR6
GNB1L
ARMT1
Suppl. Figure S4. Results of DCMS, window, and point F_ST_ analyses of UGC against a set of closely related breeds (Kalmyk, Chianina, Romagnola, and Mongolian). Red lines indicate q-value = 0.05 for DCMS and F_ST_ = 0.4 for windows F_ST_ analyses.

### Slide 5
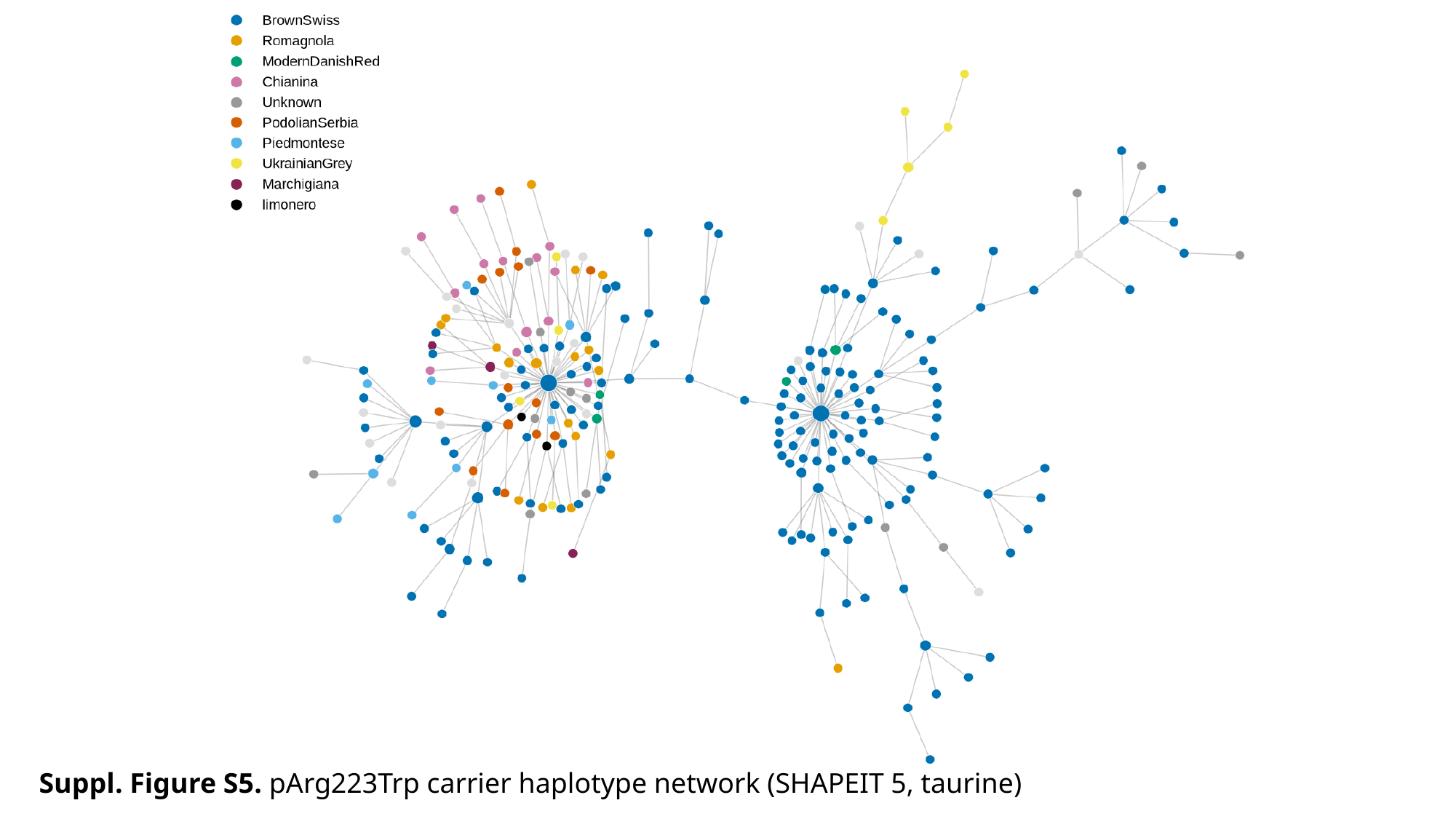

Suppl. Figure S5. pArg223Trp carrier haplotype network (SHAPEIT 5, taurine)

### Slide 6
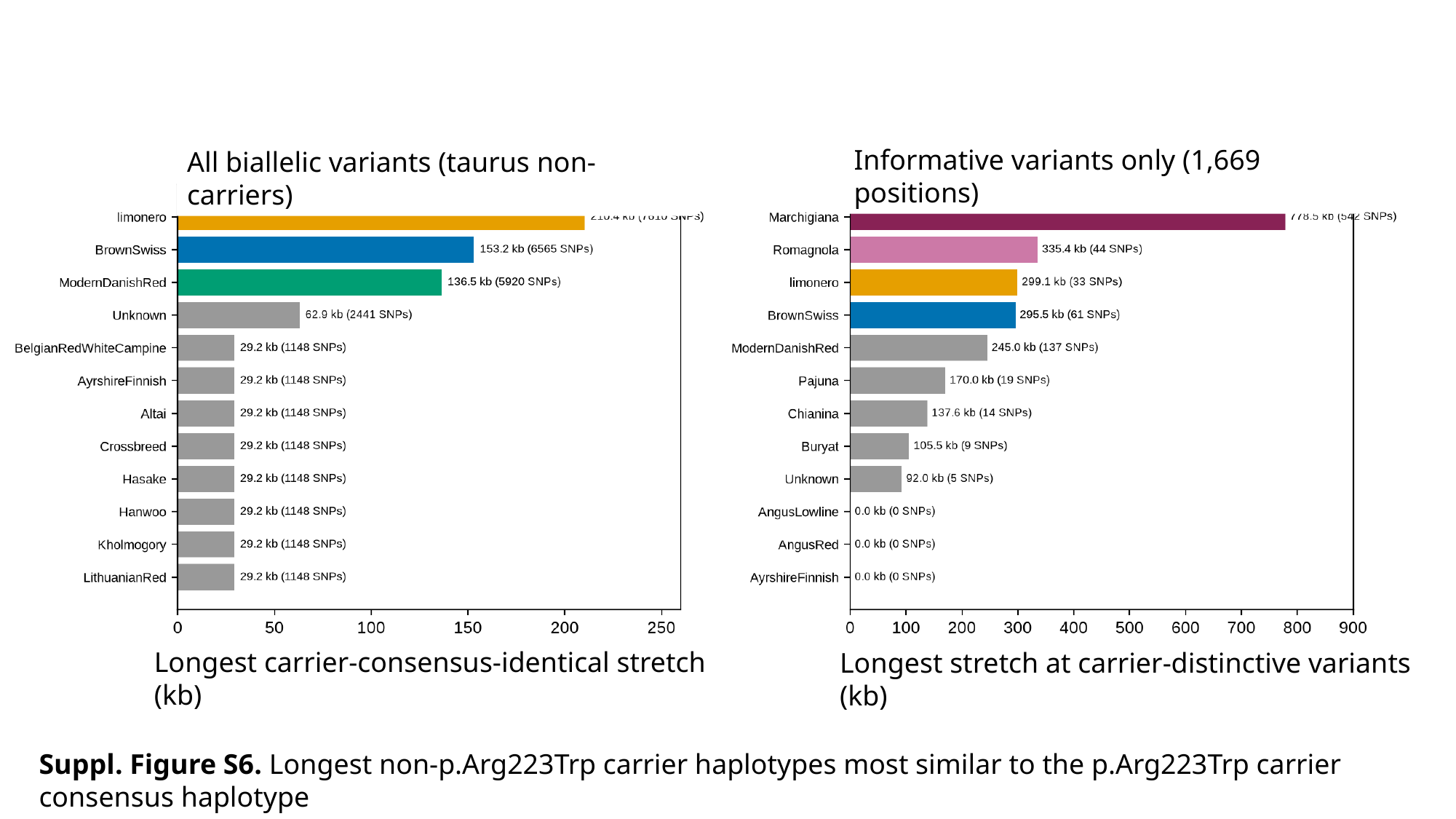

Informative variants only (1,669 positions)
All biallelic variants (taurus non-carriers)
Longest carrier-consensus-identical stretch (kb)
Longest stretch at carrier-distinctive variants (kb)
Suppl. Figure S6. Longest non-p.Arg223Trp carrier haplotypes most similar to the p.Arg223Trp carrier consensus haplotype

### Slide 7
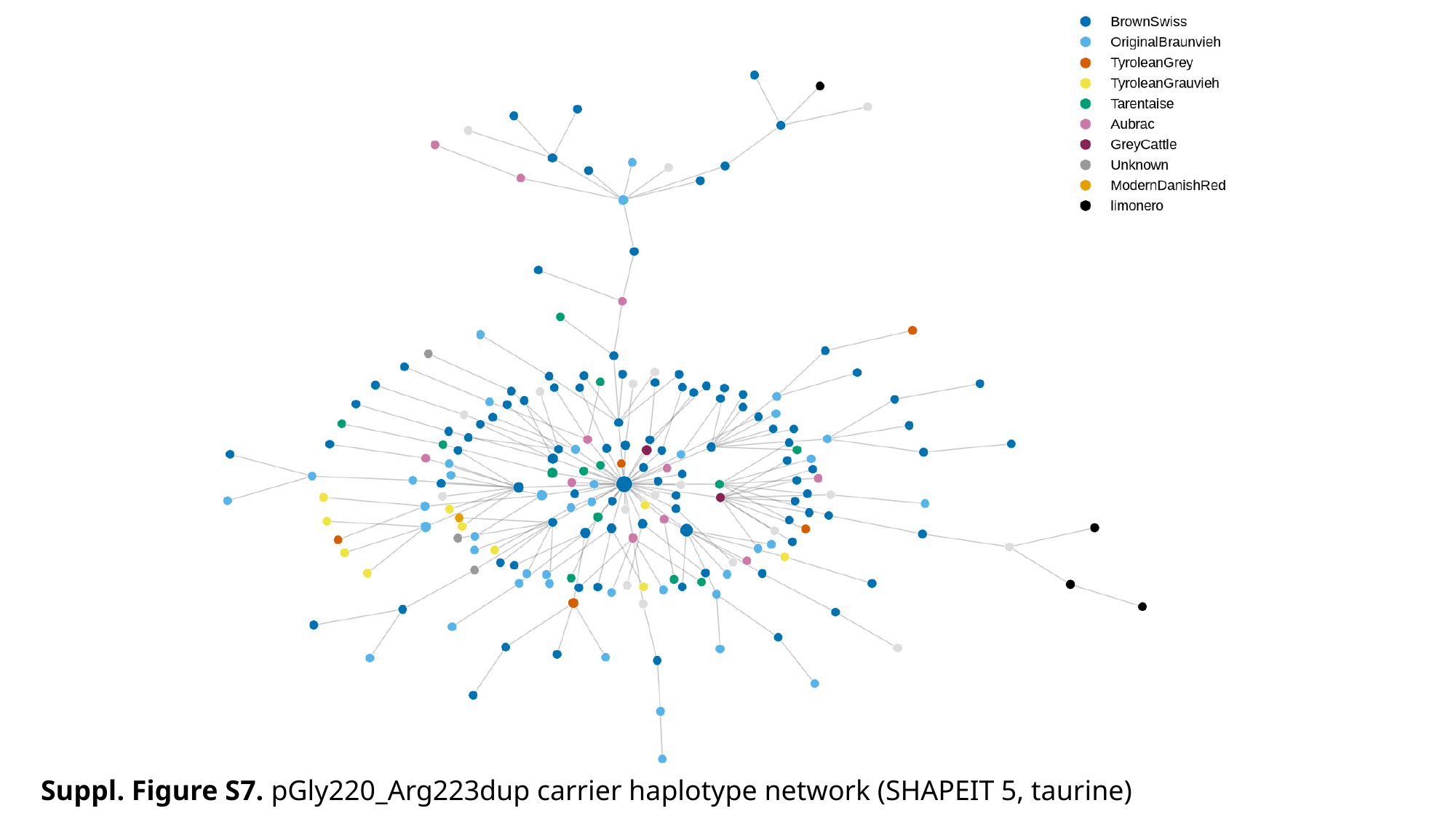

Suppl. Figure S7. pGly220_Arg223dup carrier haplotype network (SHAPEIT 5, taurine)

### Slide 8
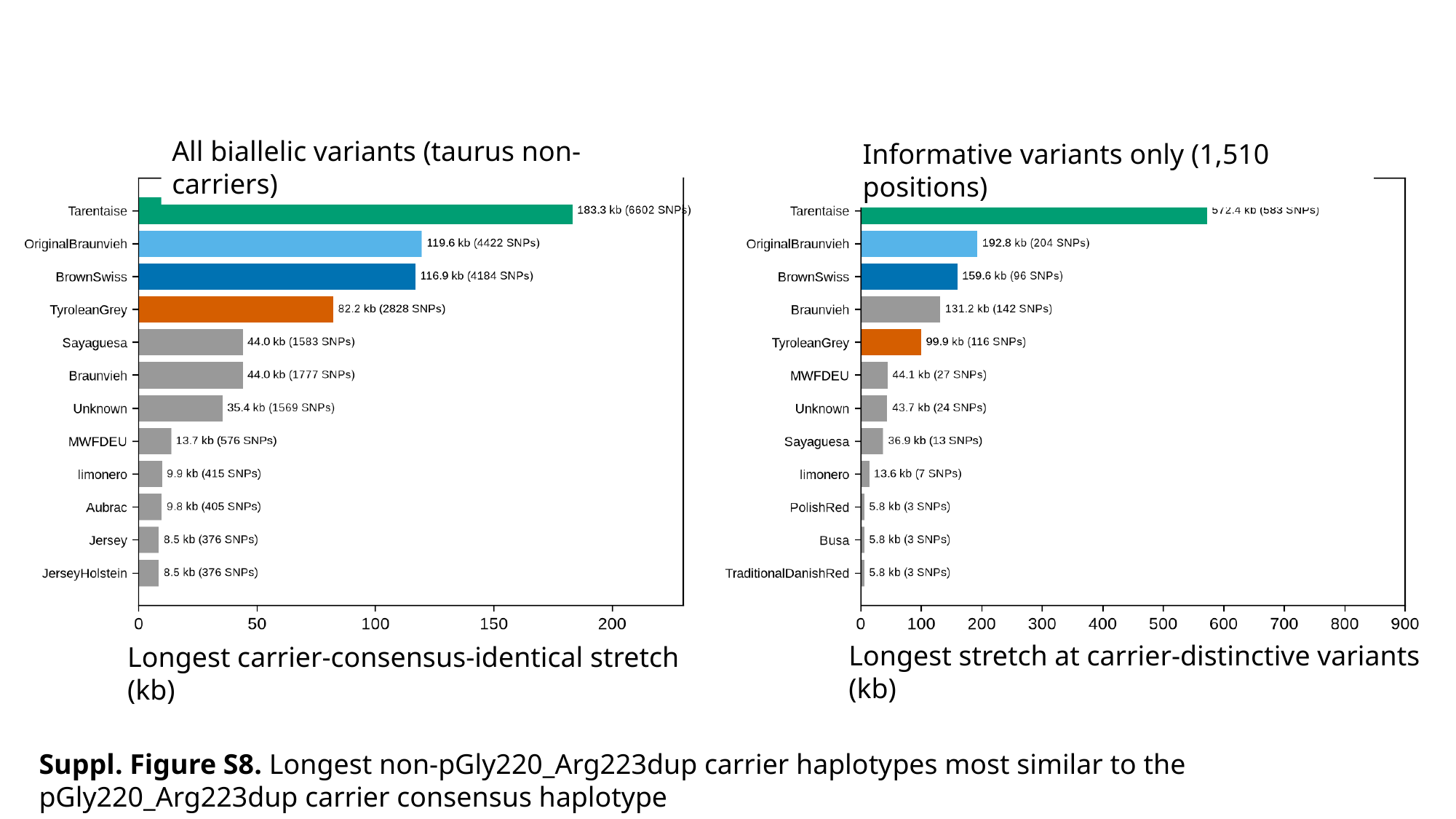

All biallelic variants (taurus non-carriers)
Informative variants only (1,510 positions)
Longest stretch at carrier-distinctive variants (kb)
Longest carrier-consensus-identical stretch (kb)
Suppl. Figure S8. Longest non-pGly220_Arg223dup carrier haplotypes most similar to the pGly220_Arg223dup carrier consensus haplotype
